# Xist RNA Dependent and Independent Mechanisms Regulate Dynamic X Chromosome Inactivation in B Lymphocytes

**DOI:** 10.1101/2025.01.27.635124

**Authors:** Natalie E. Toothacre, Kiara L. Rodríguez-Acevedo, Keenan J. Wiggins, Christopher D. Scharer, Montserrat C. Anguera

## Abstract

X-Chromosome Inactivation (XCI) involves epigenetic pathways to equalize X-linked gene expression between female and male mammals. XCI is dynamic in female B cells, as cytological enrichment of Xist RNA and heterochromatic marks on the inactive X-chromosome (Xi) are absent in naïve B cells yet return following mitogenic stimulation. Here, we asked whether any heterochromatic histone marks are present on the Xi in naïve B cells, and whether Xist RNA is required for their deposition and retention following stimulation. We find that the Xi in naïve B cells is depleted for H2AK119Ub and H3K9me3 but enriched for DNA methylation and H3K27me3, which maintain an Xist RNA-dependent epigenetic memory of XCI. Upon stimulation, Xist-independent H3K27me3 and Xist-dependent H2AK119Ub modifications accumulate across the Xi with temporal and spatial specificity. Our findings reveal the importance of Xist RNA, H3K27me3, and H2AK119Ub marks for the epigenetic integrity of X-linked genes across the Xi following female B cell stimulation.

## INTRODUCTION

X chromosome inactivation (XCI) is an epigenetic process that equalizes X-linked gene expression between female and male eutherian mammals^1^. This process is initiated with the upregulation of the long non-coding RNA Xist from the future inactive X chromosome (Xi) resulting in transcriptional silencing^2,3^. First, active histone marks (H3K27acetyl)^4^ and RNA Polymerase II^5^ are evicted from one X chromosome, then silencing histone modifications such as H3K27me3^4,6–8^, H2AK119-ubiquitin (Ub)^4,9^, and H3K9me3^10,11^ accumulate, followed by enrichment of DNA methylation (DNAm) at CpG islands and promoter regions^12–14^. In most somatic cells, the epigenetic features of the Xi can be visualized cytologically using RNA FISH and immunofluorescence (IF), where Xist RNA appears as a ‘cloud’ structure coating the Xi and the heterochromatic histone modifications H3K27me3 and H2AK119Ub form a focus^7,9^. These epigenetic modifications transmit a memory of transcriptional silencing into daughter cells with each division.

During the formation of the Xi, Xist RNA recruits Polycomb Repressive Complexes (PRC) 1 and 2 across the chromosome with temporal specificity^8,15,16^. Following the eviction of activating marks and gene silencing^4^, the RING1A/RING1B subunits of non-canonical PRC1 deposit H2AK119Ub along the Xi independent of PRC2^17^. PRC2 subunits can recognize H2AK119Ub mark and deposit H3K27me3 marks across the Xi^18,19^. Finally, canonical PRC1 recognizes H3K27me3 and adds H2AK119Ub modifications^17^. As initiation progresses, PRC marks are most rapidly enriched at intergenic regions after which they spread into active gene bodies, promoters, and putative enhancers^4^. Although the precise dynamics of H3K9me3 deposition on to the Xi by Setdb1 during murine XCI initiation have not been reported, it has been shown that H3K27me3 and H3K9me3 are enriched in mutually exclusive chromosomal blocks along the Xi chromosome in epiblast-derived stem cells (epiSCs) and mouse embryonic fibroblasts (MEFs), for maintenance of transcriptional silencing ^10,11^.

While most genes on the Xi are transcriptionally silenced, some genes ‘escape’ inactivation and are expressed. Some escape genes, such as X-Y gene pairs encoding proteins essential for housekeeping functions, are expressed from the Xi across many cell types, while others are expressed from the Xi in a cell type specific manner^20,21^. XCI escape genes are depleted for H3K27me3 and H2AK119Ub modifications at promoters and gene bodies on the Xi^16^, and have lower levels of promoter DNAm as compared to genes subject to inactivation^22^.

In most somatic cells, XCI and transcriptional silencing is maintained by continuous localization of Xist RNA and heterochromatic marks at the Xi. For example, in MEFs, deletion of Xist RNA Repeat B reduces H3K27me3 and H2AK119Ub from the Xi^16^. Additionally, deletion of *RingA/B* (PRC1) or *Eed* (PRC2) leads to the complete loss of the respective mark, a reduction of the reciprocal mark, and Xist RNA mis-localization^16^. The interdependence between Xist RNA, PRC1, and PRC2 suggests that both the canonical and non-canonical PRC pathways are involved in XCI maintenance in female somatic cells.

Unlike most somatic cells, B cells from both humans and mice have a unique mechanism of XCI maintenance in which cytological enrichment of the canonical Xist RNA cloud and H3K27me3 and H2AK119Ub^23–25^ is not observed in naïve B cells, but is restored following antigen stimulation. Specifically, Xist RNA transcripts re-localize to the Xi by 12 hours after *in vitro* stimulation, and this accumulation continues through 24 hours, prior to the first cell division^24^. Foci for H3K27me3 and H2AK119Ub co-localize with Xist RNA ‘clouds’, detected in the Xi territory using IF at 12-24hrs post-stimulation^24^. Surprisingly, although localization of Xist RNA and heterochromatic histone marks are dependent on B cell activation, dosage compensation of the Xi in naïve B cells is similar to that observed in B cells at 12 and 24 hours post-stimulation, with only ∼8-9% of X-linked genes being expressed^26^. However, the exact mechanism regulating XCI in either naïve or activated B cells remains enigmatic.

Here, we investigate allele-specific epigenetic changes across the X chromosomes in both naïve B cells and in mitogen-stimulated B cells and determined whether Xist RNA is required for the observed changes. We find that H3K27me3 and DNAm, but not H2AK119Ub or H3K9me3, are enriched across the Xi in naïve B cells, functioning as an epigenetic imprint for transcriptional silencing that is initially dependent upon Xist expression, but does not require Xist RNA localization for its maintenance. We find that H3K27me3 and H2AK119Ub modifications accumulate across the Xi following B cell activation, with temporal and spatial specificity. Furthermore, while H2AK119Ub accumulation is dependent upon Xist RNA, accumulation of H3K27me3 is surprisingly Xist-independent. These data highlight novel mechanisms governing X-linked gene expression in B cells and thus have critical implications for understanding molecular mechanisms underlying female-biased immune responses.

## RESULTS

### H3K27me3 marks, but not H2AK119Ub or H3K9me3, are enriched on the Xi in naïve B cells

Although *Xist* is expressed in naïve B cells, the Xi lacks cytological enrichment of Xist-RNA, H3K27me3 and H2AK119Ub (**Figure 1A**)^23,24^. Yet the Xi is mostly transcriptionally silent in naïve B cells^26^ prompting us to determine which epigenetic modifications could potentially be responsible for this effect. We used an F1 hybrid mouse model of skewed XCI generated by mating female C57BL/6 mice harboring a heterozygous *Xist* deletion to male *M. m. castaneous* mice (‘F1 mus x cast’). XCI in F1 females with the inherited *Xist* deletion is restricted to the paternally-derived ‘cast’ X chromosome, resulting in skewed XCI in all tissues. Moreover, single nucleotide polymorphisms (SNPs) between the C57BL/6 and cast genomes can be used to distinguish reads originating from either the Xi or Xa (**Figure 1A**). We isolated splenic naïve CD23^+^ B cells from female F1 mice and performed allele-specific CUT&RUN for histone tail modifications associated with gene repression (H3K27me3, H2AK119Ub, H3K9me3) and active transcription (H3K27ac). We found that the Xi has significantly more H3K27me3 (blue) in large domains across the chromosome compared to the Xa (black) (**Figure 1B**). The Xi and Xa have similar levels of H2AK119Ub (purple), yet the Xi has less H3K9me3 (green) as compared to the Xa in naïve B cells (**Figure 1B**). Quantification of the total CPM normalized reads across each X confirmed that there is a significant enrichment of H3K27me3 and similar levels of H2AK119Ub on the Xi as compared to the Xa (**Figure 1C**). However, the Xa has significantly more H3K9me3 compared to the Xi (**Figure 1C**), suggesting that Xi H3K9me3 enrichment in mice may be specific to XCI initiation and early development^10,11^. In naïve B cells, enrichment of H3K27me3 and H2AK119Ub are strongly positively correlated and located in similar domains across the Xi, and H3K27me3 and H3K9me3 are negatively correlated and exhibit enrichment in distinct regions across the Xi (**Figure 1D**), similar to other studies in MEFs and EpiSCs^10,11^.

**Figure 1:**
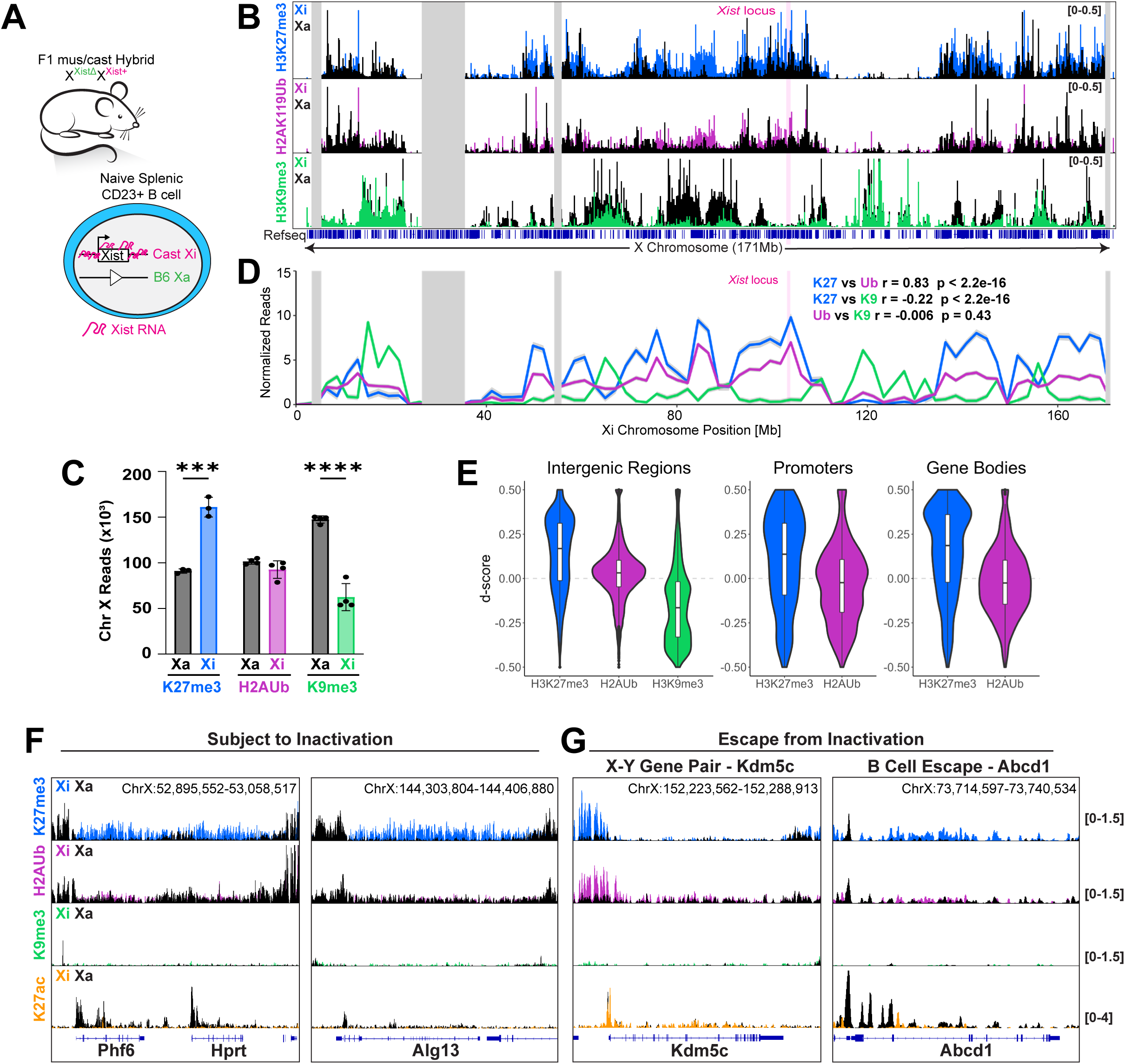
H3K27me3, but not H2AK119Ub or H3K9me3, is enriched on the Xi in naive B cells. (A) Schematic of the F1 mus x cast Hybrid model of skewed XCI. (B) Genome browser tracks of CPM normalized reads for H3K27me3 (top), H2AK119Ub (middle), and H3K9me3 (bottom) across the X chromosome. Colors (blue, purple, and green respectively) represent reads from the Xi and black represents reads from the Xa. The pink bar marks the Xist locus and gray bars indicate unmappable regions. (C) Sum of CPM normalized reads across the Xi and Xa in naive B cells. *** adj p value < 0.001, **** adj p < 0.0001 by unpaired t test. (D) Normalized read counts ±SEM in naïve B cells for H3K27me3 (blue), H2AK119Ub (purple), and H3K9me3 (green) in 10kb bins across the Xi. The pink bar marks the Xist locus and gray bars indicate unmappable regions. Pearson correlation (r) was performed between each mark. (E) D-score analysis of intergenic regions (10kb regions not overlapping genes), promoters (±1kb TSS), and gene bodies (+1kb TSS to TES). (F and G) Genome browser tracks of CPM normalized reads for H3K27me3 (blue), H2AK119Ub (purple), H3K9me3 (green), and H3K27ac (orange) at subject genes (F) and XCI escape genes (G).

To determine the genomic contexts for each heterochromatic modification, we calculated the d-score (reads^Cast^/[reads^B6^ + reads^Cast^] – 0.5) at promoter regions, gene bodies, and intergenic regions. Intergenic regions, defined as 10kb windows that do not overlap genes and their promoters (1kb upstream of the transcription start site (TSS) to the transcription end site (TES)), were enriched for H3K27me3, but not enriched for H2AK119Ub, and depleted for H3K9me3 as compared to the Xa (**Figure 1E**). Promoter regions (±1kb TSS) and gene bodies (+1kb TSS to TES) were similarly enriched for H3K27me3 but not H2AK119Ub (**Figure 1E**), indicating that H3K27me3 is present uniformly across the Xi. Gene body H3K27me3 enrichment is observed at silent genes on the Xi (*Phf6*, *Hprt*, *Alg13*) (**Figure 1F, S1I**). H3K9me3 was not quantified at promoter regions and gene bodies because it was not enriched at these sites (**Figure 1F** - green).

Next, we examined enrichment of histone modifications at repressed and escape genes on the Xi. Using CPM normalized read counts, we generated enrichment heatmaps for heterochromatic (H3K27me3, H2AK119Ub, H3K9me3) and active (H3K27ac) histone marks ±5kb of all diploid expressed X-linked genes in naïve B cells (RPKM > 1, n = 309) separated by Xi expression status (‘escape’ or ‘silent’) (**Figure S1E**)^26^. In contrast to Xi silent genes, the Xi escape genes *Kdm5c, Kdm6c,* and *Abcd1* are significantly depleted for H3K27me3 in gene bodies and downstream from the TES (**Figures 1G, S1F, S1J**; blue). H2AK119Ub was not significantly different between escape and silent genes on the Xi, except for *Kdm6a* where H3K27ac levels were also high across the gene body (**Figure S1E**, **S1G, S1J**). The B cell gene *Btk* escapes XCI in naïve B cells and has low levels of both H3K27me3 and H2AK119Ub with high levels of H3K27ac (**Figure S1J**), which is intriguing because increased *Btk* expression is a feature of autoimmune disease^27,28^. However, although not statistically significant, both H3K27me3 and H2AK119Ub marks trend towards enhanced enrichment at upstream regions of XCI escape genes, including expressed X-Y genes pairs (**Figures 1G**, **S1E** and **S1J**). This upstream enrichment is correlated with H3K27ac marks at the TSS (**Figures 1G**, **S1E** and **S1H**), suggesting that H3K27me3/H2AK119Ub may function as a barrier to prevent activation of nearby genes. Thus, the Xi in naïve B cells is enriched for H3K27me3 and not H2AK119Ub or H3K9me3 modifications.

### H2AK119Ub and H3K27me3 marks accumulate rapidly across the Xi upon B cell stimulation

As we had previously found cytological enrichment of H3K27me3 and H2AK119Ub following B cell activation *in vitro*,^23–25^ we next determined how CpG stimulation affects enrichment of silent and active histone modifications across the Xi (**Figure 2A - model**). We observed a significant accumulation of both H2AK119Ub and H3K27me3 (**Figures 2B** and **S2A**) across the Xi after stimulation. However, there was no change of H3K9me3 levels for Xi and Xa following stimulation (**Figure 2B**), suggesting that this modification may not play a regulatory role in XCI maintenance in B cells. At 24 hours post-stimulation, H2AK119Ub and H3K27me3 enrichment levels on the Xi were similar, but much higher than that observed on the Xa, which did not exhibit stimulation-dependent changes in either mark (**Figure 2B**). In 24h stimulated B cells, H3K27me3 and H2AK119Ub continue to be highly correlated with each other across the Xi, and H3K9me3 is negatively correlated with both marks (**Figure 2C**).

**Figure 2:**
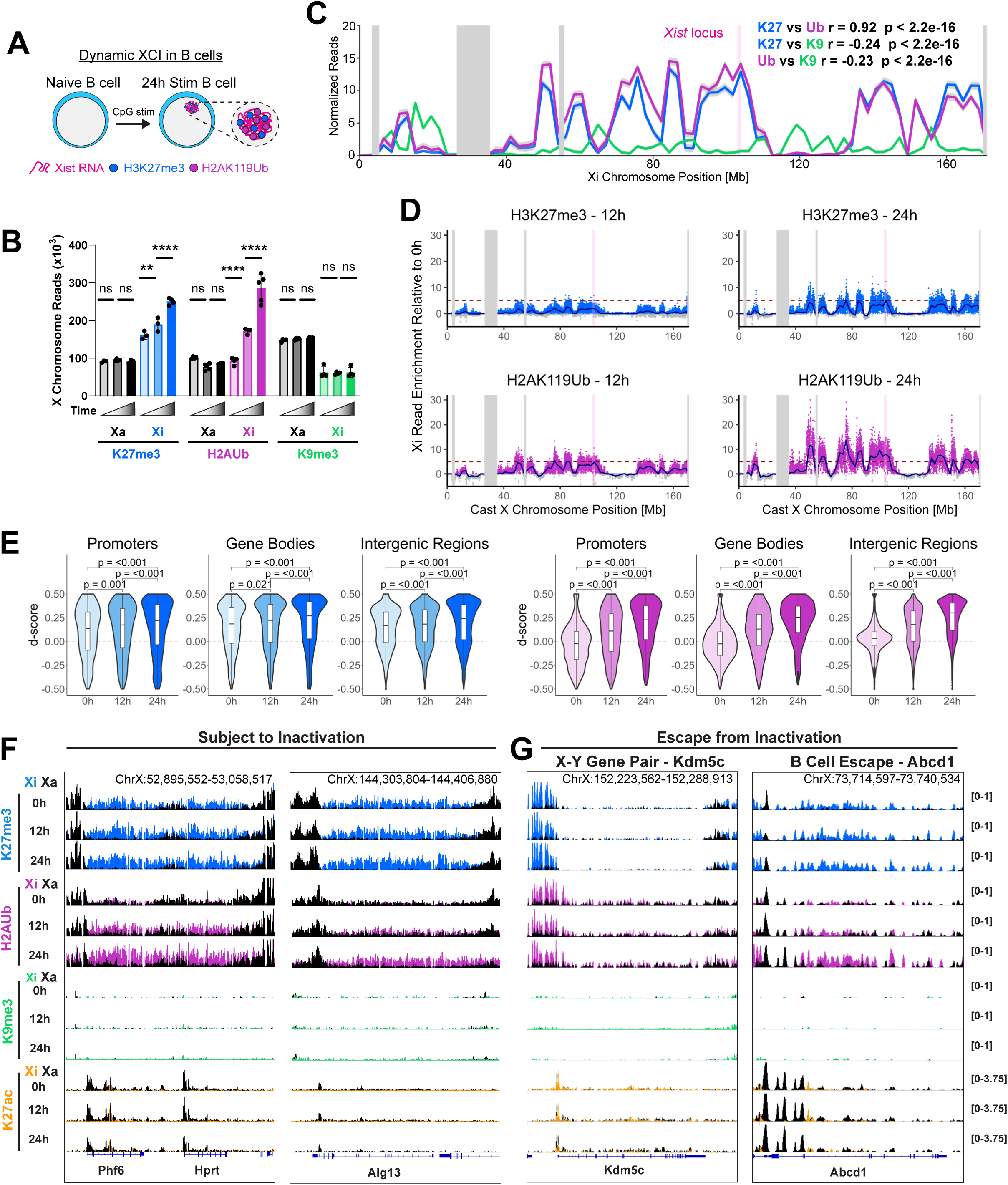
B cell activation stimulates accumulation of H3K27me3 and H2AK119Ub, but not H3K9me3 across the Xi. (A) Schematic of dynamic XCI in naïve and stimulated B cells (B) Sum of CPM normalized reads across the Xi and Xa in 0h naive, 12h stimulated, and 24h stimulated B cells. P-values calculated using one-way ANOVA followed by Holm-Sidak’s multiple comparison test, **p<0.01, ****p<0.0001 (C) Normalized 24h stim read counts for H3K27me3 (blue), H2AK119Ub (purple), and H3K9me3 (green) in 10kb bins across the Xi. The pink bar marks the Xist locus and gray bars indicate unmappable regions. Pearson correlation (r) was performed between each mark. (D) Delta accumulation of H3K27me3 (blue) and H2AK119Ub (purple) in 10kb bins (dots) across the Xi by 12h and 24h of B cell stimulation. The blue line represents the locally estimated scatterplot smoothing (LOESS) regression across all 10kb bins. The pink bar marks the Xist locus. The horizontal dotted line was drawn at 5 as a point of reference to highlight mark accumulation. (E) D-score analysis of H3K27me3 and H2AK119Ub at promoters, gene bodies, and intergenic regions. Adjusted p-values calculated using a Kruskal-Wallis test followed by a Dunn’s test with Benjamini-Hochberg correction for pairwise comparisons. (F and G) Genome browser tracks at silenced genes Phf6, Hprt, Alg13 (F) and escape genes Kdm5c, Abcd1 (G) for H3K27me3, H2AK119Ub, and H3K9me3 at 0h, 12h and 24h. Xa is overlayed in black.

We visualized the accumulation of H3K27me3 and H2AK119Ub across the Xi following B cell stimulation by calculating delta normalized read counts in 10kb bins between the 12h and 24h samples and the 0h naïve timepoint. There is substantial accumulation of both H3K27me3 and H2AK119Ub across the Xi at 12 hours of stimulation, and both modifications continue to rapidly accumulate at the same regions through 24 hours (**Figure 2D**). We quantified the accumulation of H3K27me3 and H2AK119Ub at promoters, gene bodies, and intergenic regions across the Xi using d-score analyses and observed similar rates of accumulation between these regions for both marks (**Figure 2E**). At the repressed genes *Phl6, Hprt*, and *Alg13*, there is visible accumulation of H2AK119Ub (and to a lesser extent H3K27me3), yet low levels of H3K9me3 and H3K27ac on the Xi (**Figure 2F**). For the XCI escape gene *Kdm5c*, there is continuous enrichment of both H3K27me3 and H2AK119Ub at the upstream region, and a peak for H3K27ac at the promoter, yet both H3K27me3 and H2AK119Ub are reduced across the gene body (**Figure 2G**). In contrast, we observe stimulation-dependent enrichment for H3K27me3 and H2AK119Ub, both upstream and at the gene body region for another XCI escape gene *Abcd1* (**Figure 2G**). Together, these data indicate that while silenced genes accumulate H2AK119Ub (and to a lesser extent H3K27me3), the epigenetic environment of Xi expressed genes demonstrates more variability.

Next, we examined Xi histone modification enrichment for diploid expressed X-linked genes (Xi+Xa) grouped by Xi expression status (‘escape’ or ‘silent’) (n = 314 diploid expressed genes at 24 hours). We see that at 24 hours of stimulation, H3K27me3 and H2AK119Ub have accumulated at both escape and silenced genes (**Figure S2E** – blue, purple). However, for escape genes, there is a significantly lower level of both H3K27me3 and H2AK119Ub marks along the gene body and downstream of the TES (**Figures S2E**, **S2F**, and **S2G**). Although there is still a trend towards higher enrichment of these modifications upstream of the TSS at escape genes, this difference is not statistically significant. Interestingly, H3K27ac peak intensity at escape gene promoters decreased in 24h stimulated B cells (**Figure S2H**), which may be due to the rapid accumulation of silencing marks across the Xi, leading to an indirect loss of activating marks. Thus, B cell stimulation results in significant enrichment of H2AK119Ub and H3K27me3 modifications, but not H3K9me3, at the Xi and XCI escape genes exhibit diversity of H3K27me3 and H2AK119Ub enrichment patterns in naïve and stimulated B cells.

### DNAm is enriched at CpG islands and promoters across the Xi with B cell stimulation induced DNAm changes at some regions

Because the Xi in naïve B cells is dosage compensated^26^ and contains H3K27me3, we asked whether DNAm, an epigenetic modification strongly associated with gene repression, was present across the Xi. We performed allele-specific whole genome bisulfite sequencing (WGBS) in naïve and 24 hour stimulated B cells. For naïve B cells, we found that promoter regions and CpG Islands (CGIs) are highly methylated on the Xi (∼76% and 67% respectively) as compared to the Xa (**Figures 3A** and **3B**). Gene body DNAm, associated with active transcription^29,30^, is ∼89% on the Xi compared to ∼95% for the Xa (**Figure 3C**). Similarly, in 24h stimulated B cells, we see ∼76%, 64%, and 89% DNAm at promoters, CGIs, and gene bodies on the Xi, respectively (**Figures 3D-F**). When we partition genes by expression status, we see that XCI escape genes have a significantly lower level of promoter DNAm in naïve B cells and trending to lower levels of promoter DNAm in 24h stimulated B cells (**Figures S3A-C**).

**Figure 3:**
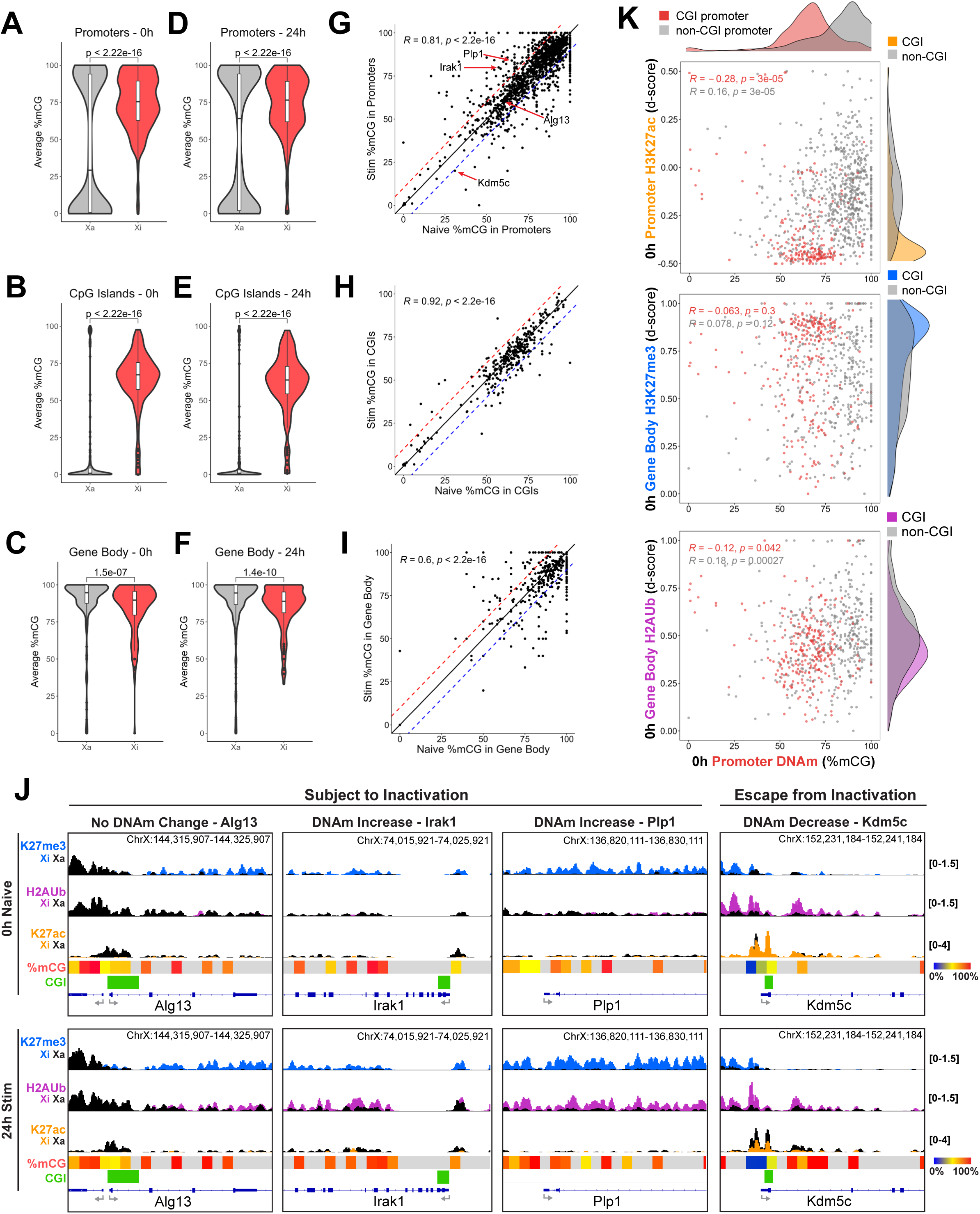
Elevated DNA methylation at promoters and CGIs on the Xi correlate with H3K27me3 enrichment in B cells. (A) Percent DNA methylation at promoter regions (±1kb TSS) on the Xa and Xi in 0h naive B cells. (B) Percent DNA methylation at CpG Islands on the Xa and Xi in 0h naive B cells. (C) Percent DNA methylation at gene bodies (+1kb TSS to TES) on the Xa and Xi in 0h naive B cells. (D-H) As A-C, respectively, in 24h stimulated B cells. All P-values calculated using Wilcoxon rank sum test. (G-I) Correlation between 0h and 24h percent DNA methylation at promoters (G), CpG Islands (H), and gene bodies (I). Red dotted line indicates an increase of 10% in DNA methylation during stimulation. Blue dotted line indicates a decrease of 10% in DNA methylation during stimulation. Pearson correlation (r) between 0h and 24h is reported. (J) 10kb windows at genes that display varying DNA methylation dynamics during stimulation. (K) Correlation between percent DNA methylation at promoter regions and promoter H3K27ac, gene body H3K27me3, or gene body H2AK119Ub in 0h naïve B cells, partitioned by presence of a CpG island within the promoter region. Pearson correlation (r) is reported.

Next, we asked if DNAm levels changed at specific promoters, CGIs, and gene bodies after B cell stimulation (24h), which is before the first B cell division. We correlated the percentage of DNAm at these 3 regions comparing naïve cells and 24h stimulated cells and see a strong correlation in DNAm between 0h and 24h, indicating that DNAm levels are largely constant (**Figures 3G-I**). Surprisingly, we see that DNAm is variable in promoters, CGIs, and gene body regions, with specific examples varying by more than 10% in stimulated B cells (**Figures 3G-I**, above and below dashed lines). For example, the *Irak1* promoter contains a CGI that exhibits decreased DNAm; the *Plp1* promoter has increased DNAm; and the CGI within the *Kdm5c* promoter has decreased DNAm (**Figure 3J**). Thus, B cell activation stimulates chromosome-wide epigenetic remodeling of both histone tail modifications and DNAm across the Xi.

Because the Xi exhibits strong correlation between DNAm with heterochromatic histone marks in female somatic cells^31^, we investigated whether X-linked promoter DNAm in naïve B cells correlates with histone modifications. We partitioned X-linked genes by CGI association (CGI promoter = CGI overlapping ±1kb of TSS) and found that genes with CGI promoters have a lower %DNAm (∼67%) as compared to non-CGI promoters (∼86%) (**Figures 3K** and **S3D**). Interestingly, X-linked genes with CGI promoters are depleted for activating promoter H3K27ac and enriched for silencing gene body H3K27me3 (**Figure 3K**), suggesting that the H3K27me3 imprint in naïve B cells is preferentially retained at intermediately methylated CGI-associated X-linked genes, likely because PRC2 targets unmethylated CG dense regions^32,33^. We also see a trend for CGI promoter genes having lower levels of gene body H2AK119Ub (**Figure 3K**). Thus, transcriptional silencing on the Xi in naïve B cells is maintained through retention of both H3K27me3 and DNAm imprints which maintain a ‘memory’ transcriptional repression in the absence of Xist RNA localization.

### H2AK119Ub and H3K27me3 show distinct temporal and spatial enrichment dynamics across the Xi with B cell stimulation

During Xi formation, H2AK119Ub marks are added prior to H3K27me3 deposition, and both heterochromatic marks accumulate at intergenic regions and later at silenced genes^4^. Thus, we asked whether there is temporal and spatial specificity for enrichment of H2AK119Ub across the Xi following B cell stimulation. We quantified gene body d-scores at X-linked genes with sufficient read coverage, selected genes that accumulate H2AK119Ub, and performed k-means clustering to generate 3 temporal clusters specific for the Xi – early, steady, and late accumulation (**Figure 4A**). We identified a cluster of Xi-linked genes with “early” H2AK119Ub accumulation, where the majority of the modification is deposited between 0h and 12h of stimulation (n = 262 genes), and a cluster with “late” H2AK119Ub accumulation, deposited between 12h and 24h (n = 248 genes) (**Figure 4A**). Early accumulation of H2AK119Ub occurs at regions close to the *Xist* locus, further from XCI escape genes, and in gene poor, LINE rich, and SINE poor regions (**Figure 4B**). Early and late H2AK119Ub accumulation groups are equally enriched for genes with CGI promoters (**Figure 4C**).

**Figure 4:**
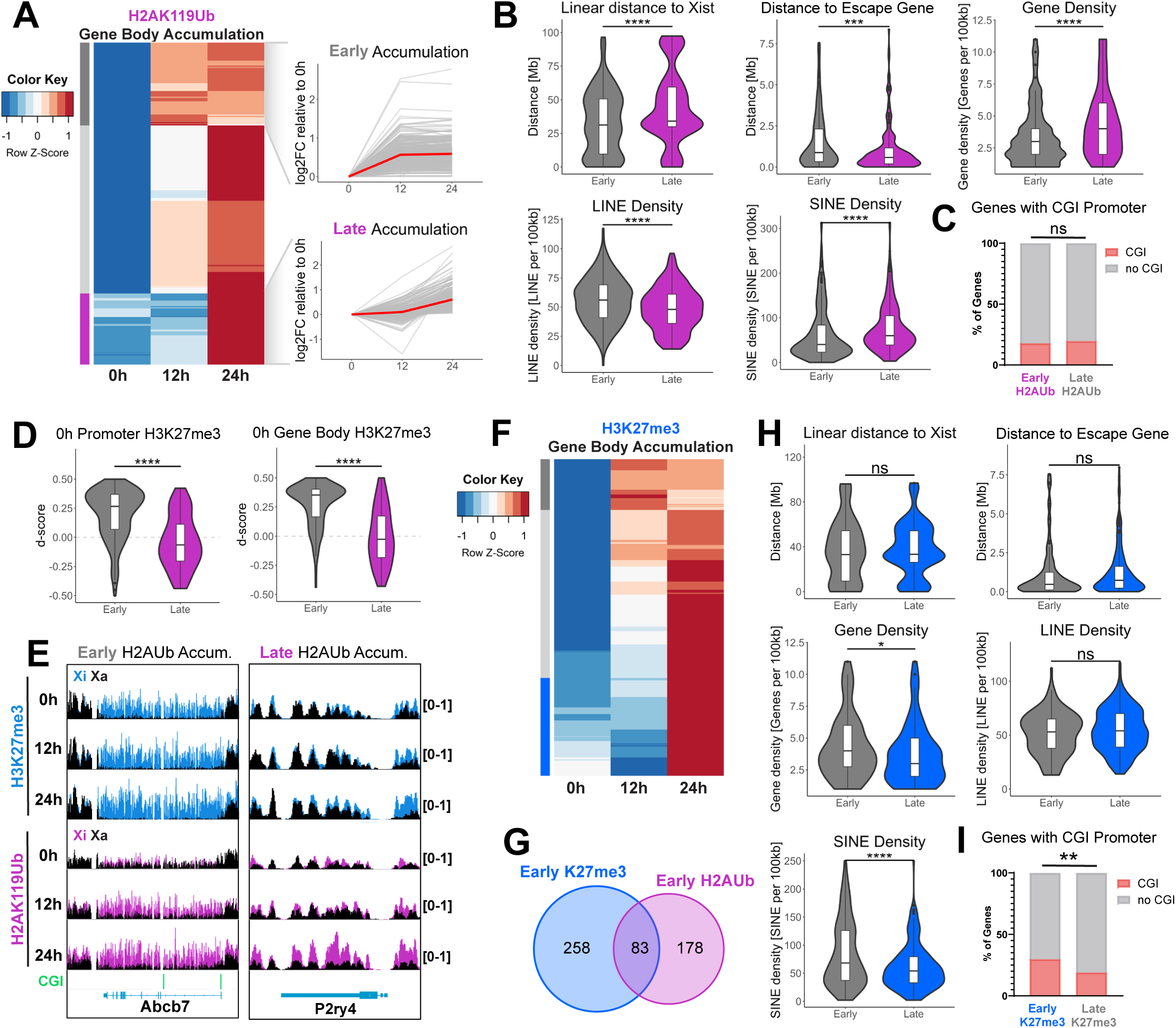
Temporal analysis of H2AK119Ub and H3K27me3 reveals preferential accumulation at specific regions on the Xi. (A) Accumulation dynamics of H2AK119Ub at genes that gain enrichment during the stimulation time course. (left) Heatmap of H2AK119Ub gene body d-scores at 0h, 12h and 24h post stimulation sorted using k-means unsupervised clustering, (right) log2 fold change of gene body d-scores across the time course for each temporal cluster. Each gray line represents one gene and the red line represents the median log2 fold change of the cluster. n = 290 for Early Accumulation, n = 248 for Late Accumulation. (B) Quantification of genomic features associated with Early and Late H2AK119Ub accumulation. Shown are linear distance to Xist, linear distance to an escape gene, gene density, LINE density, and SINE density. *** adj p < 0.001, **** p < 0.0001 by pairwise Wilcox test. (C) Percent of genes in Early and Late H2AK119Ub Accumulation groups with CGI promoters. p-value calculated using two-sided chi-square test. (D) Quantification of naive B cell H3K27me3 at genes with Early and Late H2AK119Ub accumulation. **** p < 0.0001 by pairwise Wilcox test. (E) Gene browser tracks of CPM normalized reads on the Xa (black) and Xi for H3K27me3 (blue) and H2AK119Ub (purple) at 0h, and 12h and 24h post stimulation. Shown is a gene with Early H2AK119Ub enrichment (left) and a gene with Late H2AK119Ub enrichment (right). (F) Accumulation dynamics of H3K27me3 at genes that gain enrichment during the stimulation time course. Heatmap of H3K27me3 gene body d-scores at 0h, 12h and 24h post stimulation sorted using k-means unsupervised clustering. n = 341 for Early Accumulation, n = 247 for Late Accumulation. (G) Overlap between H2AK119Ub Early Accumulation and H3K27me3 Early Accumulation groups. (H) Quantification of genomic features associated with Early and Late H3K27me3 Accumulation. Shown are linear distance to Xist, linear distance to an escape gene, gene density, LINE density, and SINE density. *** adj p < 0.001, **** p < 0.0001 by pairwise Wilcox test. (I) Percent of genes in Early and Late H3K27me3 Accumulation groups with CGI promoters. ** p < 0.01 by two-sided chi-square test.

Fascinatingly, we find that X-linked genes with early H2AK119Ub accumulation are enriched for H3K27me3 at their promoters and gene bodies in naïve B cells, and genes that have late H2AK119Ub accumulation are not enriched for H3K27me3 (**Figures 4D**, **S4A** and **S4B**). These accumulation dynamics are evident at the early accumulation gene *Abcb7* and the late accumulation gene *P2ry4* (**Figure 4E**). This observation suggests that canonical PRC1 may be able to quickly target genes pre-marked with H3K27me3 at 0h, leading to the rapid accumulation of H2AK119Ub, whereas non-canonical PRC1 may be required to place the pioneering silencing mark at genes not enriched for H3K27me3 at 0h leading to a slower enrichment of both marks.

Next, we repeated the temporal analysis for H3K27me3 accumulation dynamics and identified 341 Xi-linked genes with early H3K27me3 enrichment (dark grey bar) and 247 X-linked genes with late H3K27me3 enrichment (**Figure 4F**, blue bar). There is minimal overlap between early H2AK119Ub enriched genes and early H3K27me3 enriched genes, suggesting that PRC1 and PRC2 may be acting across the Xi independently (**Figure 4G**). While there is no correlation between H3K27me3 accumulation and linear distance to the *Xist* locus, XCI escape genes, or LINE density, early H3K27me3 enrichment occurs in gene rich and SINE rich regions which is at odds with early H2AK119Ub enrichment (**Figure 4H**). Additionally, there is a greater proportion of genes with CGI promoters in the early H3K27me3 accumulation group (**Figure 4I**). Altogether, this data demonstrates that H2AK119Ub and H3K27me3 are preferentially accumulating at distinct regions across the Xi during B cell stimulation.

To investigate PRC1 and PRC2 spreading dynamics on the Xi following B cell stimulation, we performed allele-specific CUT&RUN for Ring1B (PRC1 subunit) and Suz12 (PRC2 subunit) in naïve, and 12h and 24h stimulated B cells. We observe robust Ring1B and Suz12 enrichment peaks at the *Hox* locus on chromosome 6 and expected increases in binding profiles with B cell stimulation (**Figures S4D**, **S4F**, and **S4G**). However, Xi-wide correlational analysis of Ring1B occupancy with H2AK119Ub enrichment (**Figure S4H**) and Suz12 occupancy with H3K27me3 enrichment (**Figure S4I**) revealed that the Xi is very lowly enriched for both Ring1b and Suz12, consistent with previous observations that did not see an enrichment of Ezh2 (a PRC2 subunit) on the Xi in MEFs^15^. Xi Suz12 and Ring1B peaks were only able to be detected at regions of high H3K27me3 and H2AK119Ub enrichment, such as at the XCI escape gene *Eif2s3x* (**Figure S4E**). We did not observe a difference with Ring1B enrichment at ±5kb of X-linked genes in early and late H2AK119Ub groups (**Figure S4J**). While we are unable to determine the spreading dynamics of PRC1 and PRC2 across the Xi with certainty, we observe that H3K27me3 and H2AK119Ub display distinct spatial and temporal accumulation across the Xi following B cell stimulation.

### Xist RNA deletion, but not PRC2 chemical or genetic perturbation, alters the H3K27me3 and H2AK119Ub environment across the X chromosome in naïve B cells

To determine if H3K27me3 levels across the Xi in naïve B cells depended on PRC2, we next used chemical inhibition^34,35^ or genetic deletion^36^ approaches. We injected female F1 mus x cast mice daily for 7 days with the PRC2 inhibitor EPZ6438^34^ and then isolated splenic CD23+ naïve B cells and CD23- splenocytes (**Figures S5A**). We found that global H3K27me3 levels did not change in CD23+ naïve B cells, yet there were visible reductions of H3K27me3 in CD23- splenocytes at days 6-7 (**Figure S5B**). CUT&RUN experiments for H3K27me3 in B cells from treated F1 mus x cast mice confirmed that H3K27me3 levels did not change on either autosomes or the X-chromosomes in PRC2 inhibitor vs DMSO treated B cells (**Figure S5C-D**). Finally, the impact of genetic deletion of *Ezh2* splenic CD23+ naïve B cells from female ‘Ezh2 cKO’ (Ezh2^-/-^; CD19-Cre+) mice as assessed by Western blot analyses revealed slight reductions with global H3K27me3 levels (**Figure S5F**) while CUT&RUN analyses revealed no differences with H3K27me3 enrichment on autosomes and X-chromosomes (**Figure S5G-H**). Thus, chemical and genetic inhibition of PRC2 does not reduce H3K27me3 levels in naïve B cells.

As *Xist* deletion in somatic cells results in reductions of both H3K27me3 and H2AK119Ub levels on the Xi,^16^ we next determined if deletion of *Xist* would impact the epigenetic integrity of the Xi in B cells. To delete *Xist* in naïve splenic B cells, we crossed *Xist*^fl/fl^ mice to *Mb1*^+/Cre^ mice, then generated homozygous Xist cKO/cKO (‘Xist cKO’) female offspring through intercross matings (**Figure 5A**). The Mb1-Cre deletion deletes exons 1-3 of *Xist* in the B cell lineage starting at the pro-B cell stage^37^ thus all mature B cells lack Xist expression^26,38^ (**Figures 5A** and **S5I**). We isolated splenic CD23^+^ B cells from female WT (Xist^+/+^; Mb1^+/cre^ or Xist^fl/fl^; Mb1^+/+^) and Xist cKO mice and performed CUT&RUN for H3K27me3, H2AK119Ub, and H3K27ac modifications. We are unable to determine allele-specificity as WT and Xist cKO mice are on the C57BL/6 background with random XCI, therefore X chromosome reads are combined and originate from both the Xa and Xi.

**Figure 5:**
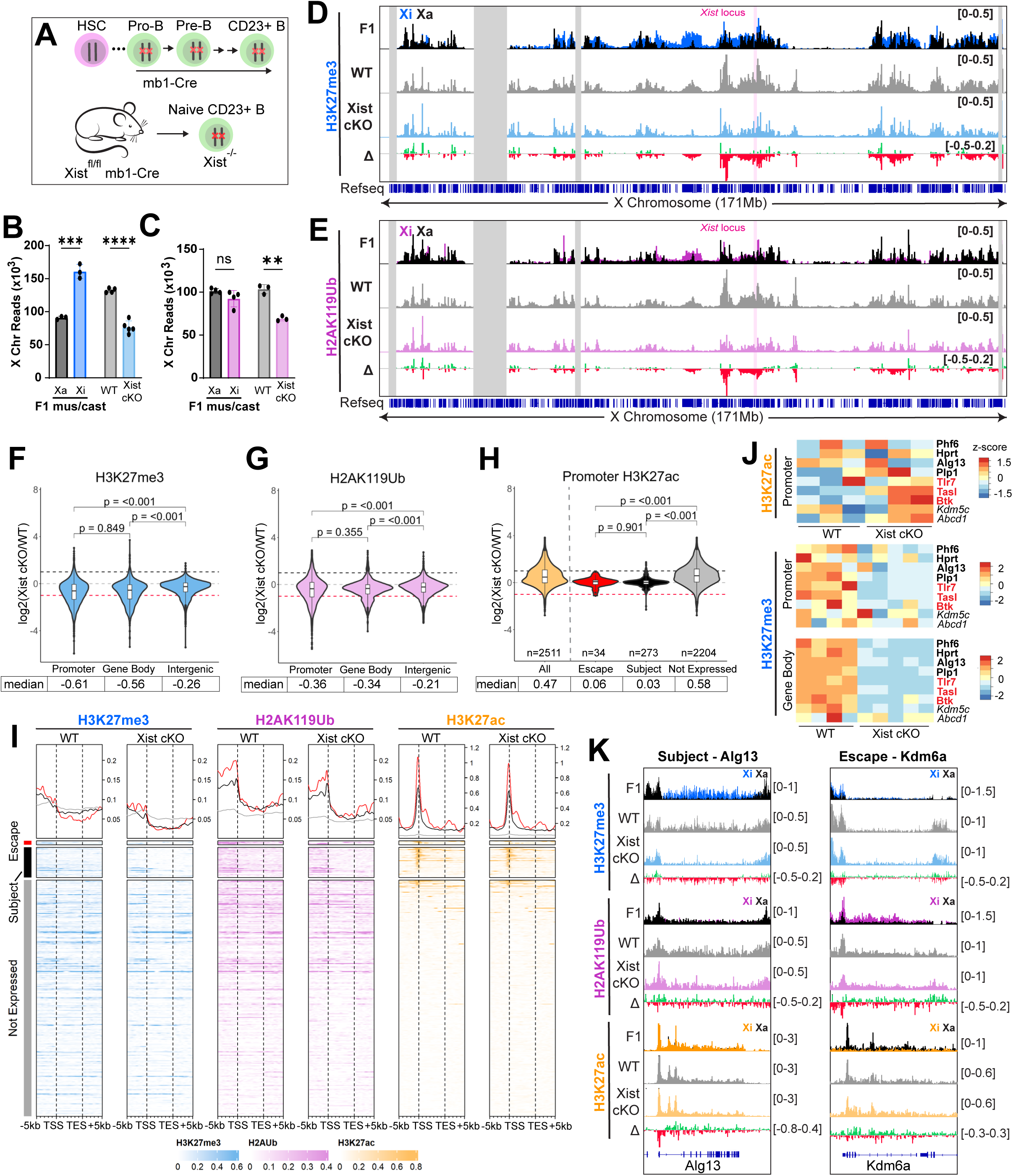
*Xist* expression is required to maintain H3K27me3 and H2AK119Ub marks on the X in naive B cells. (A) Schematic of Xist RNA deletion in naive CD23^+^ B cells with mb1-Cre. (B and C) Sum of CPM normalized H3K27me3 (B) and H2AK119Ub (C) reads on the X chromosome in F1 mus x cast (Xi and Xa), WT (mb1-cre+), and Xist cKO (Xist^fl/fl^; mb1-cre+) mice. ** adj p value < 0.01, *** adj p value < 0.001, **** adj p < 0.0001 by unpaired t test. (D and E) Genome browser tracks of CPM normalized reads for H3K27me3 (D) and H2AK119Ub (E) across the X chromosome F1 mus x cast (Xi and Xa), WT, and Xist cKO mice. Δ track represents the difference between Xist cKO and WT. (F and G) H3K27me3 (F) and H2AK119Ub (G) log2(cKO/WT) at promoters, gene bodies, and intergenic regions. (H) H3K27ac log2(Xist cKO/WT) at promoters for all genes on the X chromosome (left), then partitioned into expression status (escape, subject, and not expressed). Adjusted p-values calculated using a Kruskal-Wallis test followed by a Dunn’s test with Benjamini-Hochberg correction for pairwise comparisons. (I) Enrichment heatmap of H3K27me3 (blue), H2AK119Ub (purple), and H327ac (orange) ±5kb of all X-linked genes. (J) Changes in promoter (H3K27ac and H3K27me3) and gene body (H3K27me3) levels at select genes subject to inactivation (bold), immunity-related genes important for B cell function (red), or genes escaping inactivation (italics). (K) Genome browser zoom-in at Alg13 (subject to inactivation) and Kdm6a (escape from inactivation).

We first quantified H3K27me3 and H2AK119Ub CPM normalized reads on the X chromosomes in WT and Xist cKO naïve B cells and compared them to the previously quantified levels on the Xa and Xi in F1 mice (**Figures 5B, 5C**, from **1D**). As expected, the combined Xa+Xi levels of H3K27me3 and H2AK119Ub marks in WT naïve B cells are comparable to the Xa and Xi in F1 samples (**Figures 5B** and **5C**, grey bars). Interestingly, Xist cKO samples have significantly decreased H3K27me3 and H2AK119Ub levels, comparable or less than to levels for the Xa, indicating that the Xi H3K27me3 epigenetic imprint is lost with *Xist* deletion (**Figure 5B** and **5C**). To confirm that epigenetic mark read coverage is lost specifically from the Xi in naïve B cells, we compared X chromosome-wide read enrichment patterns in WT and Xist cKO to naïve B cell F1 samples. The H3K27me3 enrichment profile in Xist cKO samples (light blue) closely resembles Xa enrichment from F1 mice (black), and subtraction between WT and cKO samples (Δ) reveals that all regions with significant loss in H3K27me3 overlap regions of Xi enrichment in F1 samples (blue) (**Figure 5D**). Similarly, loss of H2AK119Ub enrichment in Xist cKO samples as compared to WT (Δ) is localized to sites with remaining H2AK119Ub on the Xi in naïve F1 samples (**Figure 5E**), suggesting that Xist deletion selectively decreases H3K27me3 and H2AK119Ub marks from the Xi and not the Xa.

Next, we asked whether loss of H3K27me3 and H2AK119Ub marks in Xist cKO naïve B cells are localized to promoters, gene bodies, or intergenic regions. Interestingly, we see that promoters and gene bodies lose more H3K27me3 and H2AK119Ub than intergenic regions (**Figure 5F**-**G** and **S5J**-**M**). We also quantified enrichment changes of H3K27ac marks at gene promoters across the X chromosome and detected H3K27ac accumulation at X-linked promoters of genes not expressed in naïve B cells (**Figure 5H**). To determine how *Xist* deletion impacts H3K27me3, H2AK119Ub, and H3K27ac enrichment based on X-linked gene expression, we generated enrichment heatmaps ±5kb for 3 groups of X-linked genes: ‘escape’, ‘subject to XCI’ (but expressed on Xa), and ‘not expressed’ (silent on Xa and Xi) (**Figure 5I**). Xist cKO naïve B cells have the most H3K27me3 reductions at X-linked ‘subject to XCI’ genes, and both ‘subject to XCI’ and escape X-linked genes lose more gene body H3K27me3 with *Xist* deletion compared to ‘not expressed’ genes (**Figures 5I**, **S5N**, and **S5O**). Xist cKO naïve B cells exhibit reduced H2AK119Ub marks for escape, ‘subject to XCI’, and not expressed X-linked genes (**Figures 5I**, **S5P**, and **S5Q**). While promoter H3K27ac levels increase for the ‘not expressed’ group of X-linked genes with *Xist* deletion (**Figure 5H**), the baseline level of H3K27ac is much lower for ‘not expressed’ genes compared to the other 2 groups (**Figure 5I**), suggesting that the increased H3K27ac are unlikely to result in gene reactivation.

Because *Xist* deletion in naïve B cells results in upregulated expression of just 2 X-linked genes (*Cfp, Ftx*)^26^, we asked whether *Xist* deletion would affect enrichment of epigenetic modifications at 6 X-linked immunity related genes with roles in autoimmune disease (*Btk, Tasl, Tlr7*), compared to some XCI escape genes (Kdm5c, Abcd1) and known ‘subject to XCI’ genes (*Phf6, Hprt, Alg13, Plp1*). We found that Xist cKO naïve B cells have reduced H3K27me3 levels compared to WT samples at gene bodies and promoters for these 9 X-linked genes, yet there is no change with H3K27ac levels in Xist cKO B cells (**Figure 5J**). Xist deletion changes H3K27me3 enrichment across the ‘subject to XCI’ gene (*Alg13*) and an XCI escape gene (*Kdm6a*), with minimal impact on H2AK119Ub and H3K27ac (**Figure 5K**). There is reduced H3K27me3, but not H2AK119Ub or H3K27ac, levels at gene bodies and promoters for other X-linked immunity related genes in Xist cKO naïve B cells (**Figure S5R**). We correlated X-linked gene expression with changes to epigenetic histone mark occupancy and observe increased enrichment of H3K27ac at X-linked promoters of ‘not expressed’ (grey dots) and some ‘subject to XCI’ genes (black dots) in Xist cKO naïve B cells, yet no significant correlation between gene expression changes and H3K27ac levels (**Figure S5S**). We also observed reduced levels of H3K27me3 across most X-linked gene bodies and promoters of XCI escape, ‘subject to XCI’ and ‘not expressed’ genes in Xist cKO B cells, except for *Cfp* (**Figure S5T** and **U**), which exhibits higher expression in Xist cKO naïve B cells, suggesting that reduced H3K27me3 does not alter X-linked gene expression immediately following *in vitro* stimulation.

### Xist RNA and H3K27me3 marks are required for accumulation of both H2AK119Ub and H3K27me3 on the Xi in stimulated B cells

To determine how *Xist* deletion, and reduced X-linked H3K27me3 occupancy impacts recruitment of H2AK119Ub across the Xi following B cell stimulation, we isolated and stimulated B cells from Xist cKO and WT female mice for CUT&RUN analyses of H3K27me3, H2AK119Ub, and H3K27ac occupancy (**Figure 6A**). As expected, Xist RNA was exclusively detected in female WT B cells but not in Xist cKO B cells (**Figure 6B**). We quantified accumulation of X-linked CPM normalized reads with stimulation (24h/0h) and observe significant reductions in H3K27me3 accumulation (**Figure 6C** and **S6A**) and H2AK119Ub accumulation (**Figure 6D** and **S6B**) in Xist cKO B cells compared to WT cells. In contrast, *Xist* deletion in B cells does not change H3K27me3 or H2AK119Ub enrichment on Chromosome 13 (**Figures S6C-E**). Across the length of the X chromosome, *Xist* deletion prevents accumulation of H3K27me3 at the PRC enriched domains (grey regions for WT) and there is a decrease in H2AK119Ub chromosome-wide (**Figures 6E** and **6F**). To examine whether the Xist cKO accumulation defects are localized to promoters, gene bodies, or intergenic regions, we calculated the 0 to 24 hour log2-fold change in normalized read counts for WT and cKO stimulated B cell samples. While WT samples gain H3K27me3 at all regions, in Xist cKO samples promoter H3K27me3 does not change with stimulation (median log2FC = 0.02) and H3K27me3 is lost from gene bodies and intergenic regions (median log2FC = −0.15 and −0.20 respectively) (**Figure 6G**). *Xist* deletion impairs stimulation induced H3K27ac accumulation at X-linked promoters (**Figure S6F**). X-linked CGI promoters are enriched for H3K27me3 in naïve B cells, yet *Xist* deletion prevents H3K27me3 accumulation at both CGI promoters and non-CGI promoters, indicating that either Xist RNA itself or H3K27me3 imprints are required for accumulation following stimulation (**Figure 6H**). *Xist* deletion also reduced H2AK119Ub accumulation at promoters, gene bodies, and intergenic regions after stimulation (**Figure 6I**). Finally, we asked how early and late H2AK119Ub accumulation genes (identified in **Figure 4**) are impacted by *Xist* deletion and loss of the H3K27me3 imprints. Early accumulation genes are enriched for H3K27me3 in WT naïve B cells, and *Xist* deletion results in greater impairment in H2AK119Ub accumulation at early genes than at late genes (median log2FC = −0.22 and −0.15, respectively) (**Figure 6J**). For example, X-linked *Abcb7* is an ‘early accumulation’ gene with a CGI promoter, and *Xist* cKO B cells exhibit reduced H3K27me3 and H2AK119Ub levels (**Figure 6K**, left). The X-linked gene *P2ry4* is a ‘late accumulation’ gene with a non-GCI promoter, and Xist cKO stimulated B cells have reduced H2AK119Ub levels (**Figure 6K**, right). Thus, both Xist RNA and the H3K27me3 X-linked imprints are required for proper epigenetic remodeling across the Xi in stimulated B cells.

**Figure 6:**
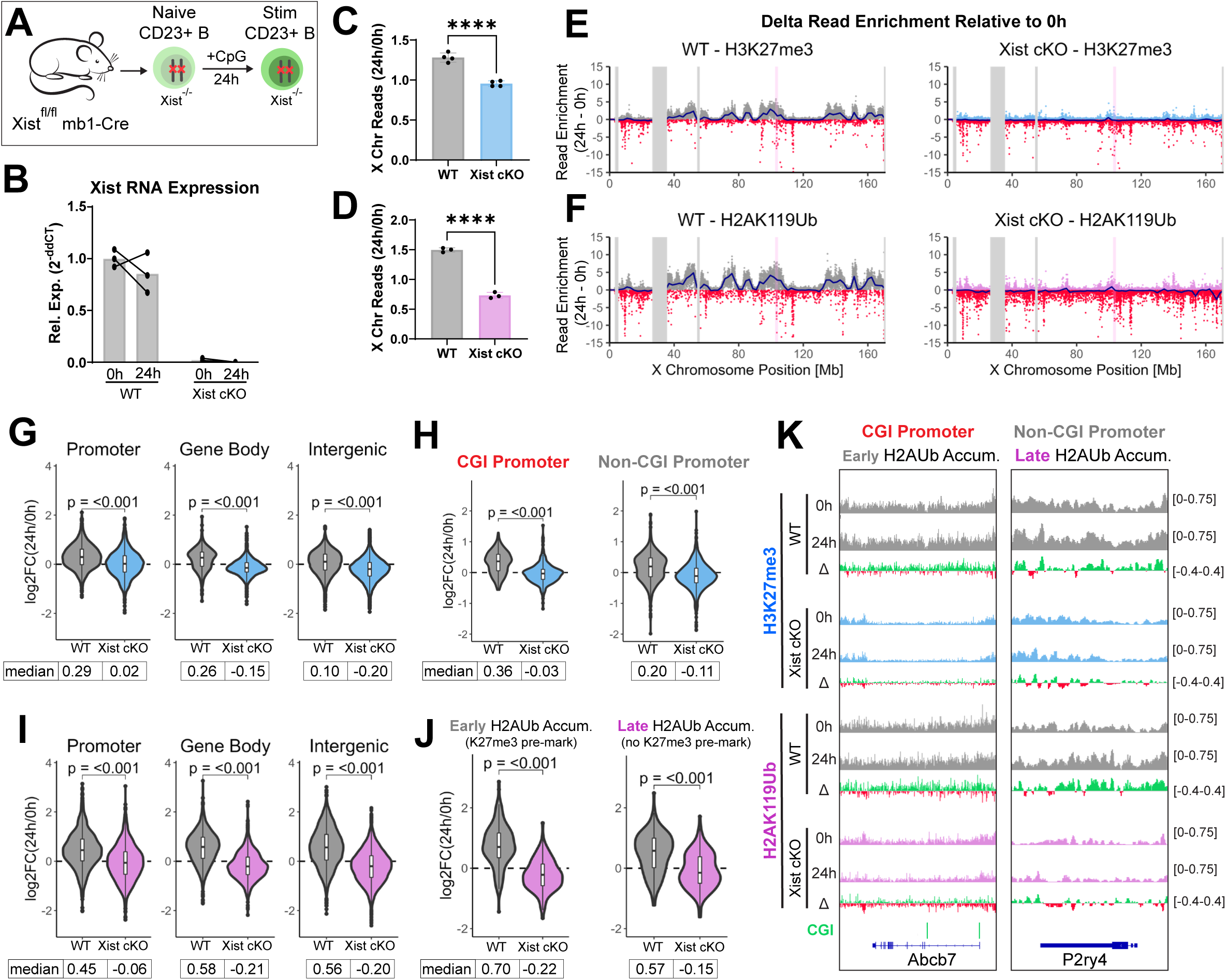
Xist RNA and H3K27me3 marks are required for H2AK119Ub and H3K27me3 enrichment on the X following B cell stimulation. (A) Schematic of in vitro stimulation of Xist cKO CD23^+^ B cells. (B) Xist RNA expression in 0h naive and 24h stimulated B cells from n= 3 WT (mb1-cre+) and n=3 Xist cKO (Xist^fl/fl^ mb1-cre+) mice. (C and D) Fold change in total CPM normalized X-linked reads between 0h naïve and 24 hour stimulated WT and Xist cKO B cells for H3K27me3 (C) or H2AK119Ub (D). **** p < 0.0001 by unpaired t-test. (E and F) Delta read enrichment (24h - 0h) of H3K27me3 (E) and H2AK119Ub (F) in 10kb bins across the X chromosome in WT (left) and Xist cKO (right) B cells. The blue line represents the locally estimated scatterplot smoothing (LOESS) regression across all 10kb bins. The pink bar marks the Xist locus. (G) H3K27me3 log2(24h/0h) at promoters, gene bodies, and intergenic regions in WT and Xist cKO B cells. (H) H3K27me3 log2(24h/0h) at gene bodies (TSS to TES) at genes with CGI promoter or non-CGI promoters in WT and Xist cKO B cells. (I) H2AK119Ub log2(24h/0h) at promoters, gene bodies, and intergenic regions in WT and Xist cKO B cells. (J) H2AK119Ub accumulation (log2(24h/0h)) at genes with “Early Accumulation” (left) and “Late Accumulation” (right) of H2AK119Ub in WT and Xist cKO B cells. P-values calculated by pairwise Wilcox test. (K) Gene browser tracks of CPM normalized reads in WT (gray) or Xist cKO (light blue or light purple) samples at 0h and 24h post stimulation. Δ = 24h - 0h. Shown is a gene with a CGI promoter and Early H2AK119Ub enrichment (left) and a gene with no CGI promoter and Late H2AK119Ub enrichment (right).

### X-linked H3K27me3 imprints in naïve B cells are sufficient for accumulation of H3K27me3, but not H2AK119Ub, in stimulated B cells

As *Xist* deletion *in vivo* may induce selective pressures that indirectly impact XCI, we next asked whether *Xist* deletion *ex vivo*, immediately preceding B cell stimulation, would impact stimulation-induced accumulation of H3K27me3 and H2AK119Ub across the X chromosome. We isolated splenic naïve CD23^+^ B cells from Xist^fl/fl^ mice (where *Xist* is intact), performed an *ex vivo* deletion with TAT-Cre recombinase protein (‘TAT KO’)^39,40^, and stimulated the cells for 24 hours with CpG (**Figure 7A**). Xist RNA levels are decreased by ∼65% at 24h of stimulation in female TAT KO B cells (**Figure 7B**), and because the half-life of Xist RNA is 1.9 hours in CD23^+^ B cells^24^, Xist RNA levels are mostly depleted before robust re-localization to the Xi occurs. We performed CUT&RUN for H3K27me3, H2AK119Ub, and H3K27ac in WT (Xist^+/+^ TAT-Cre treated) and TAT KO (Xist^fl/fl^ TAT-Cre treated) samples at the naïve and 24h stimulated timepoints.

**Figure 7:**
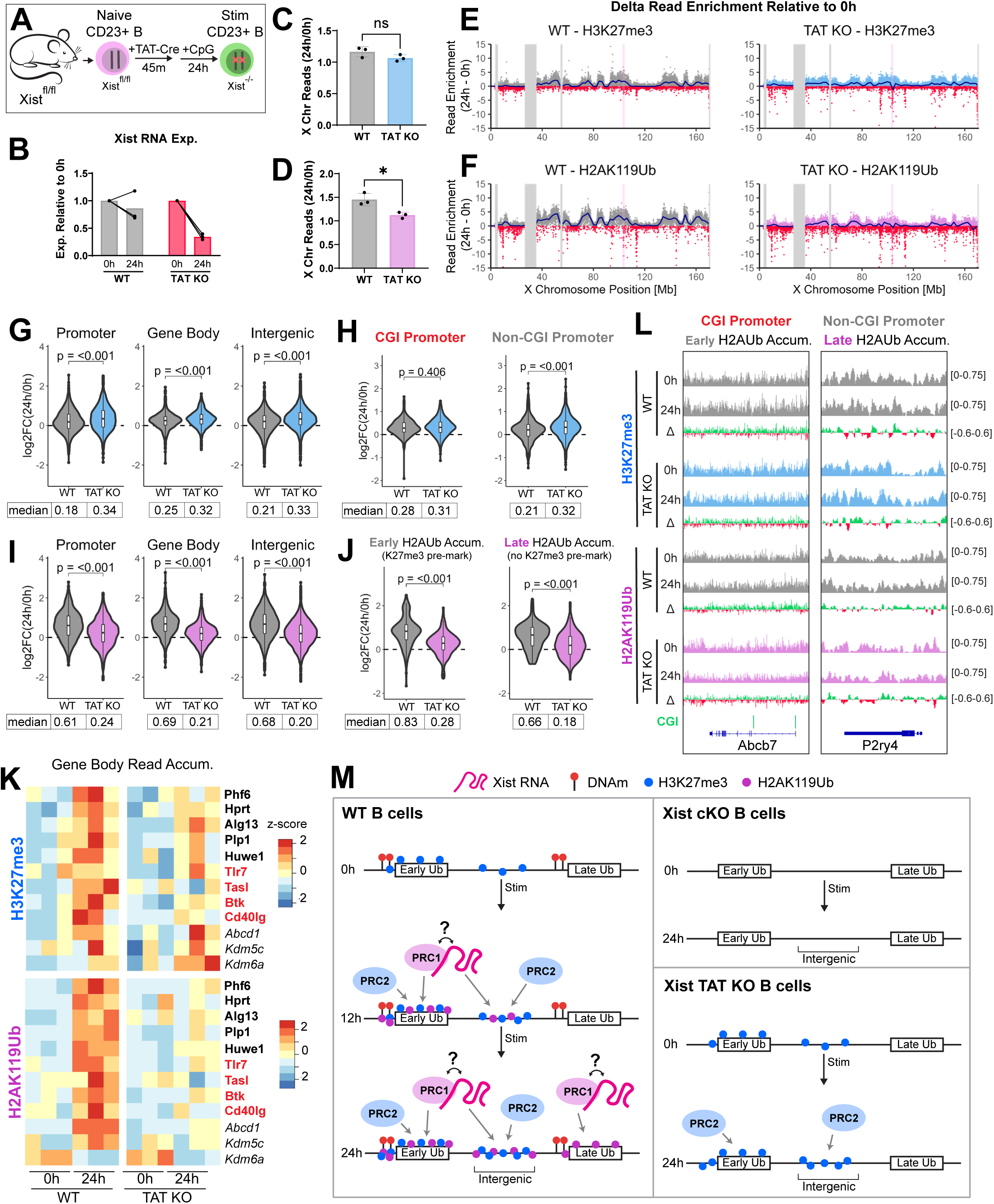
*Xist* deletion prior to B cell activation prevents H2AK119Ub, but not H3K27me3, accumulation on the X. (A) Schematic representation of *ex vivo* Xist deletion using TAT-Cre recombinant protein. (B) Xist RNA expression in 0h naive and 24h stimulated B cells treated *ex vivo* with TAT-Cre recombinant protein, n= 3 WT and n=3 TAT KO (Xist^fl/fl^). (C and D) Fold change in CPM normalized X-linked reads between 0h naive and 24h stimulated WT and KO B cells for H3K27me3 (C) and H2AK119Ub (D) * p < 0.05 by unpaired t-test. (E and F) Delta read enrichment (24h - 0h) of H3K27me3 (E) and H2AK119Ub (F) in 10kb bins across the X chromosome in WT (left) and cKO (right) B cells. The blue line represents the locally estimated scatterplot smoothing (LOESS) regression across all 10kb bins. The pink bar marks the Xist locus. (G) H3K27me3 log2(24h/0h) at promoters, gene bodies, and intergenic regions in WT and TAT KO B cells. (H) H3K27me3 log2(24h/0h) at gene bodies (TSS to TES) at genes with CGI promoter or non-CGI promoters in WT and TAT KO B cells. (I) H2AK119Ub log2(24h/0h) at promoters, gene bodies, and intergenic regions in WT and TAT KO B cells. (J) H2AK119Ub accumulation (log2(24h/0h)) at genes with “Early Accumulation” (left) and “Late Accumulation” (right) of H2AK119Ub in WT and TAT KO B cells. P-values calculated by pairwise Wilcox test. (K) Changes in and gene body H3K27me3 and H2AK119Ub levels at select genes subject to inactivation (bold), immunity-related genes important for B cell function (red), or genes escaping inactivation (italics). (L) Gene browser tracks of CPM normalized reads in WT (gray) or TAT KO (light blue or light purple) samples at 0h and 24h post stimulation. Δ = 24h - 0h. Shown is a gene with a CGI promoter and Early H2AK119Ub enrichment (left) and a gene with no CGI promoter and Late H2AK119Ub enrichment (right). (M) Model of epigenetic remodeling of the Xi during dynamic XCI.

We quantified fold change in H3K27me3 CPM normalized reads (24h/0h) across the X chromosome during stimulation and see no significant difference in H3K27me3 accumulation when comparing WT and TAT KO B cells, suggesting that the H3K27me3 imprints in naïve B cells are sufficient for H3K27me3 accumulation despite reduced levels of Xist RNA (**Figure 7C** and **S7A**). However, H2AK119Ub accumulation on the X chromosome requires Xist RNA, as there is a significant decrease in accumulation when comparing WT and TAT KO samples (**Figures 7D** and **S7B**). Across the X chromosome, there are visibly similar levels of H3K27me3 accumulation across the PRC domains (**Figures 7E** and **S7F**) however, there are smaller peaks of H2AK119Ub accumulation after stimulation (**Figures 7F** and **S7F**). Examination of the log2-fold change in read accumulation at X-linked promoters, gene bodies, and intergenic regions revealed that TAT KO stimulated B cells exhibit relatively similar levels of H3K27me3 accumulation as WT samples (**Figure 7G**), including both CGI-associated and non-CGI promoters (**Figure 7H**).

Next, we quantified H2AK119Ub accumulation at X-linked promoters, gene bodies, and intergenic regions in TAT KO B cells, and observe significantly less accumulation compared to WT cells (**Figure 7I**). Because stimulation-induced H2AK119Ub accumulation is directed by pre-existing H3K27me3 marks, we examined H2AK119Ub accumulation in TAT KO samples at both ‘early accumulation’ X-linked genes (pre-marked with H3K27me3) or late accumulation genes (lacking H3K27me3). Interestingly, we find that H2AK119Ub marks do not accumulate at either early or late genes (**Figure 7J**). For example, at the X-linked gene *Abcb7* which is ‘early accumulation’ gene that also contains CGIs at the promoter, we observe somewhat lower H327me3 levels and reduced H2AK119Ub enrichment in TAT KO stimulated samples (**Figure 7L**; left). For the X-linked gene *P2ry4* (late accumulation and lacks CGI), there is also reduced H2AK119Ub enrichment in TAT KO stimulated cells (**Figure 7L**; right). Examination of select X-linked genes that are subject to XCI in B cells (*Phf6, Hprt, Alg13, Huwe1*- bold), immunity-related genes (*Tlr7, Tasl, Btk, Cd40lg* - red), and XCI escape genes (*Abcd1, Kdm5c, Kdm6a* - italic) in TAT KO stimulated B cells reveals variable reduced H3K27me3 levels across genes, with immunity genes having the least H3K27me3 accumulation and XCI escape genes the most H3K27me3, yet all the selected X-linked genes exhibit reduced H2AK119Ub (**Figure 7K**). Thus, Xist deletion immediately preceding B cell stimulation exhibits variable effects on X-linked gene H3K27me3 accumulation, yet H2AK119Ub is significantly reduced across the X chromosome, revealing both Xist independent and dependent mechanisms for H3K27me3 and H2AK119Ub accumulation in female stimulated B cells.

## DISCUSSION

Here, we investigated the epigenetic profile of the Xi in naïve B cells, and how the enrichment of active and silent epigenetic modifications is altered before the first cell division following their CpG-induced stimulation. Using allele-specific CUT&RUN and WGBS approaches, we discovered that the Xi in naïve B cells is enriched with H3K27me3 and DNAm yet has low levels of both H2AK119Ub and H3K9me3 marks, suggesting an epigenetic H3K27me3 and DNAm imprint for transcriptional silencing. Notably, when *Xist* is deleted during B cell development, naïve B cells lack both H3K27me3 and H2AK119Ub marks on the Xi (**Figure 7M**; top right), suggesting that these marks are initially dependent upon Xist-RNA although the H3K27me3 marks do not appear to require continued Xist RNA localization for their maintenance. *In vitro* B cell stimulation using CpG triggers accumulation of both H3K27me3 and H2AK119Ub enrichment across the Xi, where H3K27me3 marked regions (‘Early Ub’) accumulate H2AK119Ub marks first at 12 hours, and later H2AK119Ub accumulates at H3K27me3 low/absent regions (‘Late Ub’) by 24 hours (**Figure 7M**; left). Our model implies that Xist RNA recruits PRC1, likely through hnRNPK interactions^41^, and while these interactions have not been shown in primary CD23+ B cells, there is evidence for Xist recruitment of PRC1 and PRC2 in somatic cells^16,18,41,42^. Using Xist *in vivo* (Xist cKO) and *ex vivo* (Xist TAT KO) deletion approaches in 24h stimulated B cells, we observe both Xist dependent and Xist independent mechanisms that regulate H3K27me3 and H2AK119Ub accumulation across the Xi. As noted above, when *Xist* is deleted during B cell development, naïve B cells lack both H3K27me3 and H2AK119Ub marks on the Xi, and both marks fail to accumulate at 24h post-stimulation (**Figure 7M**; top right). When Xist is deleted immediately preceding B cell stimulation, the pre-existing H3K27me3 marks direct PRC2 for additional H3K27me3 accumulation yet PCR1 is not able to deposit H2AK119Ub marks on the Xi (**Figure 7M**; bottom right). Thus, our model for XCI maintenance in B cells proposes two parallel pathways for the epigenomic remodeling of the Xi following cellular stimulation.

Compared to our previous cytological experiments ^23–25^, our allele-specific CUT&RUN analyses offers improved detection of epigenetic modifications across the Xi in naïve and *in vitro* stimulated B cells. Through quantification of various heterochromatic (H3K27me3, H2AK119Ub, H3K9me3) and active (H3K27acetyl) histone tail modifications in naïve and stimulated B cells, we see similarities and differences with XCI initiation and maintenance mechanisms across cell types. First, the Xi in both naïve B and unstimulated T cells^43^ is dosage compensated, yet lacks Xist RNA localization at the Xi and has higher levels of H3K27me3 marks compared to the Xa. In addition, there are similar levels of H2AK119Ub on the Xi and Xa in both naïve B and unstimulated T cells, which suggests conservation of epigenetic maintenance pathways for the Xi across lymphocytes. Our observation of Xi specific enrichment of H3K27me3 in B and T cells underscores the distinct properties of Xi chromatin in lymphocytes compared to other somatic cells, which is likely to have consequences for gene expression and possibly Xi specific gene reactivation across cell subsets in health and disease. Second, while H3K9me3 is cytologically enriched on the Xi in some human^44^ and marsupial^45^ somatic cells, and ChIP-seq analyses indicate that H3K9me3 is enriched on the mouse Xi at intergenic regions^10,11^, the Xa has more H3K9me3 enrichment compared to the Xi in B cells, and stimulation does not alter H3K9me3 levels across the Xi. Moreover, unlike female MEFs and epiblast cells where ∼56% of H3K9me3 blocks also contain H3K27me3^11^, H3K9me3 blocks across the Xi are negatively correlated with H3K27me3 and H2AK119Ub marks in B cells. These distinct features of the Xi chromatin in B cells, and likely also T cells, may impact the fidelity of gene silencing in lymphocytes.

Using allele-specific WGBS, we discovered that the Xi in B cells also has high levels of DNAm at promoters and CpG islands, suggesting that DNAm together with H3K27me3 imprints maintain transcriptional memory of Xi silencing independent of Xist RNA tethering on the Xi in naïve B cells. Xi promoters in both naïve and stimulated B cells have the highest levels of DNAm compared to autosomes and the Xa, consistent with a dosage compensated X chromosome^46^. In naïve and stimulated B cells, Xi promoters lacking CGIs exhibit higher levels of DNAm and lower levels of H3K27me3 compared to CGI-containing promoters on the Xi. PRC2 can associate with unmethylated CG-dense regions of DNA^32,33^, thus reduced DNAm at CGI-associated promoters may facilitate PRC2 binding to the Xi for increased retention of H3K27me3 marks. It is intriguing to consider that continuous *Xist* expression from the Xi in naïve B cells may be required to maintain both H3K27me3 and DNAm across the Xi, and future studies examining allele-specific epigenomic and gene expression perturbations modulated by DNAm regulatory enzymes are necessary.

Our study reveals that the H3K27me3 imprint, both on the Xi and globally, in naïve B cells is incredibly persistent, and only *Xist* deletion successfully reduced this mark across the Xi. Treating mice with the Ezh2 inhibitor EPZ6438 did not affect global H3K27me3 levels in splenic CD23^+^ naïve B cells by Western blot, although the mark was effectively depleted in CD23^-^ splenocytes. Using CUT&RUN, which exhibits higher sensitivity than Western blotting, there was no change in naïve B cell H3K27me3 enrichment across the X chromosome. Genetic deletion of *Ezh2* using CD19-cre^36^ did not alter H3K27me3 levels on either autosomes or the X chromosomes in splenic naïve B cells, as assessed by Western blots and CUT&RUN. The persistent retention of H3K27me3 marks in naïve B cells following EPZ6438 treatment or Ezh2 deletion may be attributable to functional redundancy of Ezh1 for PRC2 mediated deposition of H3K27me3^47^, as naïve B cells have more Ezh1 than Ezh2 protein^48^. However, *Xist* expression is required to maintain H3K27me3 and H2AK119Ub modifications across the Xi in naïve B cells. In differentiating ES cells and MEFs, after the initial deposition of H2AK119Ub by the non-canonical PRC1 complex^4^, addition of H3K27me3 and H2AK119Ub by PRC2 and PRC1 also rely on Xist RNA recruitment^16^. In MEFs, the Xi depleted of both H3K27me3 and H2AK119Ub marks did not exhibit major changes with X-linked gene expression^16^, similar to *Xist* deletion in naïve and 24h stimulated B cells^26^. However, our study demonstrates that Xist cKO reduces the epigenetic integrity of the Xi in naïve B cells, with reduced H3K27me3 and H2AK119Ub marks at X-linked immunity genes like *Tasl*. These epigenetic impairments at specific immunity X-linked genes likely persist following B cell activation and differentiation, facilitating aberrant overexpression in *Xist* deleted age-associated B cells and germinal center B cells^38^. In support, some female B cell Xist cKO mice develop spontaneous lupus-like disease with advanced age, with development of specific autoantibodies characteristic of female- biased autoimmunity and kidney pathologies^38^.

B cell stimulation results in epigenetic remodeling across the Xi which is regulated by both Xist RNA dependent and independent mechanisms. *Xist* deletion during B cell development results in the depletion of H3K27me3 marks on the Xi in naïve B cells, thereby preventing accumulation of both H3K27me3 and H2AK119Ub following B cell stimulation. However, when *Xist* is deleted immediately preceding stimulation, the Xi retains H3K27me3 marks which directs PRC2 to add more H3K27me3 marks but is insufficient for PRC1-mediated accumulation of H2AK119Ub to the Xi, despite the presence of H3K27me3 imprints. While these findings suggest that PRC2-mediated accumulation is independent of Xist RNA, our *ex vivo* Xist RNA deletion system is incomplete (∼60% deletion efficiency). Thus, while our CUT&RUN analyses may be influenced by residual Xist RNA expression, it still suggests that H3K27me3 and H2AK119Ub enrichment are differentially dependent on Xist RNA. In addition, we find that PRC1- and PRC2-mediated addition and retention of H3K27me3 and H2AK119Ub on the Xi in B cells differs from that in other cell types^16,17^. During the formation of the Xi, H2AK119Ub is added first by non-canonical PRC1 before PRC2 adds H3K27me3 marks, after which both modifications are added interdependently^17^. In MEFs, Xist RNA is necessary to maintain levels of both H3K7me3 and H2AK119Ub on the Xi^16^ similar to naive B cells. However, in stimulated B cells with the H3K27me3 imprint, H3K27me3 accumulation is independent of both Xist RNA expression and pre-existing H2AK119Ub marks on the Xi, likely proceeding through EED recognition of H3K27me3 marks for PRC2-mediated maintenance of H3K27me3 levels^49–51^.

Altogether, our study has identified the kinetics for epigenomic changes across the Xi during B cell stimulation and demonstrates the requirement of Xist RNA for H3K27me3 and H2AK119Ub modifications across the chromosome. We find that B cells utilize different epigenomic pathways on the Xi compared to other somatic cells, thus warranting additional investigation to understand how these mechanisms impact X-linked gene expression. The epigenomic changes occurring across the Xi in response to B cell stimulation provide opportunities to understand X-linked immune gene pathways which become aberrantly expressed in female-biased autoimmune diseases and will inform future studies examining epigenome and gene expression changes in pathogenic B cell subsets.

## Data availability

All sequencing data generated in this study has been deposited to the NCBI GEO database. Access data using the following accession numbers: [GSE282253, GSE282254, GSE282255, GSE282256]

## ACKNOWLEDGEMENTS

We would like to thank the PennVet CHMI for their help with sequencing; L. King for help with manuscript editing; and all members from Anguera lab for helpful discussions. We thank K. Sarma, R. Bonasio, M. Bartolomei for CUT&RUN reagents and insights on data interpretation. We would like to thank S. Henikoff for pA-MNase used for CUT&RUN experiments. We would like to thank Z. Beethem for optimizing the CUT&RUN workflow. This research was supported by NIH R01 AI134834 and R01 AI168047, Lupus Research Alliance Target in Lupus grant (to M.C.A.).

## AUTHOR CONTRIBUTIONS

Conceptualization, M.C.A. N.E.T.; Methodology, M.C.A., N.E.T. ; Investigation, N.E.T., K.L.R.; Software,.; Formal Analysis, N.E.T.; Resources,.; Manuscript Writing, N.E.T. and M.C.A; Manuscript Review & Editing, M.C.A., N.E.T.; Funding Acquisition, M.C.A.

## FIGURES & FIGURE LEGENDS

**Supplementary Figure 1: Allele-specific profiling of H3K27me3, H2AK119Ub, H3K9me3, and H3K27acetyl (ac) in naïve B cells.**

(A) Genome browser tracks of CPM normalized reads for H3K27me3, H2AK119Ub, H3K9me3, and H3K27ac at the Hox gene cluster and neighboring expressed gene Skap2. (B) Genome browser tracks of CPM normalized reads for H3K27me3 (top), H2AK119Ub (middle), and H3K9me3 (bottom) across chromosome 13. Colors (blue, purple, and green respectively) represent reads from the Cast allele and black represents reads from the Black6 allele. Gray bars indicate unmappable regions. (C) Normalized read counts in naïve B cells for H3K27me3 (blue), H2AK119Ub (purple), and H3K9me3 (green) in 10kb bins across the Cast chromosome 13. Gray bars indicate unmappable regions. Pearson correlation (r) was performed between each mark. (D) D-score analysis of intergenic regions (10kb regions not overlapping genes), promoters (±1kb TSS), and gene bodies (+1kb TSS to TES). (E) Enrichment heatmap of H3K27me3 (blue), H2AK119Ub (purple), H3K9me3 (green), and H327ac (brown) ±5kb of all expressed X-linked genes in naïve B cells. (F and G) Quantification of Upstream, Gene Body, and Downstream regions shown in (E) for H3K27me3 (F) and H2AK119Ub (G). (H) Quantification of Promoter region H3K27ac shown in (E). P-value by unpaired t-test. (I and J) Genome browser tracks of CPM normalized reads for H3K27me3 (blue), H2AK119Ub (purple), H3K9me3 (green), and H3K27ac (orange) at a subject gene (I) and escape genes (J).

**Supplementary Figure 2: Allele-specific profiling of H3K27me3, H2AK119Ub, H3K9me3, and H3K27acetyl (ac) in stimulated B cells (12 and 24 hr).**

(A) Genome browser tracks of CPM normalized reads for H3K27me3, H2AK119Ub, and H3K9me3 across the X chromosome and chromosome 13 during stimulation. (B) Normalized read counts in 24 hour stimulated B cells for H3K27me3 (blue), H2AK119Ub (purple), and H3K9me3 (green) in 10kb bins across the Cast chromosome 13. Gray bars indicate unmappable regions. Pearson correlation (r) was performed between each mark. (C and D) Delta accumulation of H3K27me3 (C) and H2AK119Ub (D) in 10kb bins (dots) across the chromosome 13 by 12h and 24h of B cell stimulation. The blue line represents the locally estimated scatterplot smoothing (LOESS) regression across all 10kb bins. (E) Enrichment heatmap of H3K27me3 (blue), H2AK119Ub (purple), H3K9me3 (green), and H327ac (brown) ±5kb of all expressed X-linked genes in 24 hour stimulated B cells. (F and G) Quantification of Upstream, Gene Body, and Downstream regions shown in (E) for H3K27me3 (F) and H2AK119Ub (G). unpaired t-test. (H) Quantification of Promoter region H3K27ac shown in (E). P-values by unpaired t-test.

**Supplementary Figure 3: DNAm is enriched at X-linked gene promoters and correlates with H3K27me3 enrichment at gene bodies in B cells.**

(A) Enrichment heatmap of DNA methylation ±5kb of all expressed X-linked genes in naïve and 24 hour stimulated B cells. Blue, yellow, and red represent 0%, 50%, and 100% DNA methylation, respectively. Gray represents regions with insufficient read coverage. (B and C) Quantification of Promoter region DNA methylation from (A) in naïve (B) and 24 hour stimulated B cells (C). P-values by unpaired t-test. (D) Correlation between percent DNA methylation at promoter regions and promoter H3K27ac, gene body H3K27me3, or gene body H2AK119Ub in 24 hour stimulated B cells, partitioned by presence of a CpG island within the promoter region.

**Supplementary Figure 4: Allele-specific profiling of Ring1b and Suz12 enrichment in B cells.**

(A and B) Quantification of H3K27me3 at genes with Early (A) and Late (B) H2AK119Ub accumulation in naïve, 12 hour stimulated and 24 hour stimulated B cells. * p < 0.01, **** p < 0.0001 by Kruskal-Wallis test followed by a Dunn’s test with Benjamini-Hochberg correction for pairwise comparisons. (C) Log2 fold change of gene body H3K27me3 d-scores across the time course for the Early and Late temporal clusters. Each gray line represents one gene and the red line represents the median log2 fold change of the cluster. (D and E) Genome browser tracks of CPM normalized reads for Ring1b and Suz12 at the Hox gene cluster (D) and at the escape gene Eif2s3x (E) in naïve, 12 hour stimulated, and 24 hour stimulated B cells. (F and G) Genome-wide differentially bound peaks between naïve and 24 hour stimulated B cells for Ring1b (F) and Suz12 (G). (H) Pearson correlation (r) between Ring1b and H2AK119Ub read density on the Xi in naïve, 12 hour stimulated, and 24 hour stimulated B cells. (I) Pearson correlation (r) between Suz12 and H3K27me3 read density on the Xi in naïve, 12 hour stimulated, and 24 hour stimulated B cells. (J) Enrichment heatmap of Ring1b and Suz12 in naïve and 24 hour stimulated B cells. Shown is ±5kb of X-linked genes that accumulate H2AK119Ub, partitioned by Early, Steady, and Late Accumulation.

**Supplementary Figure 5: Chemical and genetic inhibition of Ezh2 and Xist deletion using mb1-Cre (‘Xist cKO’) in naïve B cells.**

(A) Experimental design of PRC2 chemical inhibition (PRC2i) with EPZ6438 in F1 mus x cast females. (B) Western blot for H3K27me3 in CD23+ splenic B cells and CD23- splenocytes from F1 mice treated with DMSO, 6 days of PRC2i, or 7 days of PRC2i. (C) Normalized read counts from chromosome 4, chromosome 13, and the X chromosome quantified from H3K27me3 CUT&RUN in CD23+ splenic B cells from treated F1 mice. (D) Genome browser tracks of the X chromosome from H3K27me3 CUT&RUN in CD23+ splenic B cells from treated F1 mice. (E) Experimental design of PRC2 genetic perturbation with Ezh2^fl/fl^;CD19-cre (Ezh2 cKO) mice. (F) Western blot for H3K27me3 in CD23+ splenic B cells from WT and Ezh2 cKO mice. (G) Normalized read counts from chromosome 4, chromosome 13, and the X chromosome quantified from H3K27me3 CUT&RUN in CD23+ splenic B cells from WT and Ezh2 cKO mice. (H) Genome browser tracks of the X chromosome from H3K27me3 CUT&RUN in CD23+ splenic B cells from WT and Ezh2 cKO mice. (I) qPCR of relative Xist RNA expression in naïve B cells from WT and Xiat cKO mice (n = 3 each). (J and K) log2(Xist cKO/WT) at promoters, gene bodies, and intergenic regions on chromosome 13 for H3K27me3 (J) and H2AK119Ub (K). Adjusted p-values calculated using a Kruskal-Wallis test followed by a Dunn’s test with Benjamini-Hochberg correction for pairwise comparisons. (L and M) log10(Reads per 1kb) at promoters, gene bodies, an intergenic regions on the X chromosome for H3K27me3 (L) and H2AK119Ub (M). P-values calculated by pairwise Wilcox test. (N and O) H3K27me3 log2(Xist cKO/WT) at promoters (N) and gene bodies (O) for all genes on the X chromosome (left) then partitioned into expression status (escape, subject, and not expressed). (P and Q) H2AK119Ub log2(Xist cKO/WT) at promoters (P) and gene bodies (Q) for all genes on the X chromosome (left) then partitioned into expression status (escape, subject, and not expressed). Adjusted p-values calculated using a Kruskal-Wallis test followed by a Dunn’s test with Benjamini-Hochberg correction for pairwise comparisons. (R) log2(Xist cKO/WT) at a subset of immunity-related genes for promoter H3K27ac, promoter and gene body H3K27me3, and promoter and gene body H2AK119Ub. (S-U) Correlation between change in gene expression [log2(Xist cKO/WT)] with Xist RNA cKO^26^ and change in promoter H3K27ac (S), promoter H3K27me3 (T), and gene body H3K27me3 (U) with Xist RNA cKO [log2(Xist cKO/WT)]. Escape genes are red, genes subject to inactivation are black, not expressed genes are gray.

**Supplementary Figure 6: Quantification of H3K27me3, H2AK119Ub, and H3K27ac marks in B cells from Xist cKO mice.**

(A and B) Sum of CPM normalized H3K27me3 (A) or H2AK119Ub (B) reads across the X chromosome in 0h naive and 24h stimulated WT and Xist cKO B cells. (C and D) Sum of CPM normalized H3K27me3 (C) or H2AK119Ub (D) reads across chromosome 13 in 0h naive and 24h stimulated WT and Xist cKO B cells. * p < 0.05, ** p < 0.01, calculated by paired T-test. (E) Delta read enrichment (24h - 0h) of H3K27me3 (top) and H2AK119Ub (bottom) in 10kb bins across the X chromosome in WT (left) and Xist cKO (right) B cells. The blue line represents the locally estimated scatterplot smoothing (LOESS) regression across all 10kb bins. (F) H3K27ac log2(24h/0h) at promoters in WT and Xist cKO B cells. P-values calculated by pairwise Wilcox test.

**Supplementary Figure 7: Quantification of H3K27me3, H2AK119Ub, and H3K27ac marks in B cells from Xist TAT KO mice**

(A and B) Sum of CPM normalized H3K27me3 (A) or H2AK119Ub (B) reads across the X chromosome in 0h naive and 24h stimulated WT and TAT KO B cells. (C and D) Sum of CPM normalized H3K27me3 (C) or H2AK119Ub (D) reads across chromosome 13 in 0h naive and 24h stimulated WT and TAT KO B cells. * p < 0.05, calculated by paired T-test. (E) Delta read enrichment (24h - 0h) of H3K27me3 (top) and H2AK119Ub (bottom) in 10kb bins across the X chromosome in WT (left) and TAT KO (right) B cells. The blue line represents the locally estimated scatterplot smoothing (LOESS) regression across all 10kb bins. (F) Difference in read enrichment between WT and TAT KO samples in 10kb bins across the X chromosome [TAT KO^(24h – 0h)^ – WT^(24h – 0h)^]. The blue line represents the locally estimated scatterplot smoothing (LOESS) regression across all 10kb bins. (G) H3K27ac log2(24h/0h) at promoters in WT and TAT KO B cells. P-values calculated by pairwise Wilcox test

## METHODS

### Mice

Female F1 *Mus musculus castaneus* X C57BL/6J *Xist^fl/+^* (F1 mus x cast) mice were used for all allele-specific experiments, and were generated by mating Xist^fl/fl^ female mice to male B-actin Cre (B6N.FVB-*Tmem163^Tg(ACTB-cre)2Mrt^*/CjDswJ; strain# 019099, Jackson), then Xist^fl/+^ female mice were mated to male *Mus musculus castaneus* (Cast; strain# 000928, Jackson) mice to generate female F1 mus x cast mice^26^. Female Xist cKO mice with a B cell specific deletion were used for *in vivo Xist* deletion experiments, and were generated by mating Xist^fl/fl^ mice on a C57BL/6 background to Mb1-cre mice (B6.C(Cg)-Cd79^atm1(cre)Reth^/EhobJ; Jackson Laboratory Strain 020505)^38^. Xist^+/+^; mb1- cre^+/cre^ or Xist^fl/fl^; mb1-cre^+/+^ were used as controls for these experiments. Female Xist^fl/fl^ mice on a C57BL/6 background were used for *ex vivo Xist* deletion experiments. Female Xist^+/+^ mice were used as controls for these experiments. F1 mus x cast and Xist^fl/fl^ mice were maintained at the Penn Vet animal facility, and experiments were approved by the University of Pennsylvania Institutional Animal Care and Use Committee (IACUC). Ezh2^fl/fl^;CD19-cre and CD19-cre control mice were bred and maintained at Emory University, and approval for animals was obtained through the IACUC at Emory University. Euthanasia via carbon dioxide was used for animal sacrifice prior to isolations.

### *In vivo* PRC2 Inhibition

Female F1 *mus x cast* mice received daily 350 mg/kg of EPZ6438 (300mM EPZ6438 in 100% DMSO; AstaTech Cat # 40834) or DMSO control (100% DMSO) for 6 (n = 1, EPZ6438) or 7 (n = 1, EPZ6438; n = 1, DMSO control) days by intraperitoneal (IP) injection, and euthanized 3 hours following the final administration. Spleens were collected and CD23^+^-B cells and CD23^-^ splenocytes were isolated as detailed below.

### B cell isolation and stimulation

Splenic B cells were isolated from female F1 *mus x cast*, Xist^fl/fl^; mb1-cre^+/cre^, Xist^fl/fl^ and control mice (2-6 months) as previously described^23,24^. Spleens from Ezh2^fl/fl^;CD19-cre and control mice were shipped overnight on ice from Emory University then used for B cell isolations. Briefly, spleens were crushed to produce single cell suspensions, splenocytes were incubated with Biotin anti-mouse CD23 Antibody (Cat #101604, Biolegend) then streptavidin microbeads (Cat #130048101, Miltenyi), followed by running through a magnetized LS column (Cat #130042401, Miltenyi) to positively enrich for B cells. For naïve timepoints, B cells were immediately processed following isolation for CUT&RUN or WGBS experiments. For stimulated timepoints, B cells were cultured at 37°C in RPMI 1640 (Invitrogen) containing 10% fetal bovine serum (FBS, Gemini), 0.1% β- mercaptoethanol (Invitrogen), 1% nonessential amino acids (Invitrogen), 10mM HEPES (Invitrogen), 2mM L-glutamine (Invitrogen), 1% OPI media supplement (O5003-1VL, Sigma-Aldrich), and 1% penicillin- streptomycin (Invitrogen) (B cell media). B cells were stimulated with 1µM CpG (tlrl-1826, Invivogen) and collected after 12 hrs or 24 hrs. For TAT-Cre treatment, 10 million naïve B cells were treated with 80ug of TAT-Cre recombinase protein (Cat #EG-1001. Excellgen) in 1mL of Opti-MEM media (Cat #31985062, Thermo Fisher) for 45 minutes at 37°C. Treatment was stopped by adding 100uL of FBS and 3mL of B cell media, then cells were stimulated for 24 hours at 37°C with B cell media containing 1µM CpG.

### CUT&RUN

CUT&RUN for H3K27me3, H2AK119Ub, H3K9me3, H3K27ac, Ring1b, and Suz12 was performed using previously published methods^52^. In brief, 10uL/reaction of concanavalin A-coated beads (Cat #86057-3, Polysciences) were equilibrated with Binding Buffer (20mM HEPES, 10mM KCl, 1mM CaCl2, 1mM MnCl2) then resuspended in Wash Buffer (20mM HEPES, 150mM NaCl, 0.5mM Spermidine, 1x PIC (Cat #11697498001, Roche)). Approximately 1-1.5 million B cells were immobilized on beads in Wash Buffer by rotating for 1 hour at room temperature. Beads were resuspended in Antibody Buffer (20mM HEPES, 150mM NaCl, 0.5mM Spermidine, 1x PIC, 0.05% Digitonin, 2mM EDTA). Primary antibody for H3K27me3 (3 µg; Cat #39055, Active Motif), H3K9me3 (3 µg; Cat # ab8898, Abcam), H3K27ac (3 µg; Cat #39134, Active Motif), H2AK119Ub (∼1.5 µg; Cat # 8240S, Cell Signaling), IgG (3 µg; Cat #A01008, GenScript), Ring1b (3:50 dilution; Cat #5694, Cell Signaling Technology), or Suz12 (3:50 dilution; Cat #3737, Cell Signaling Technology) were added and beads were rotated at 4°C for 5 hr. Beads were washed with and resuspended in Dig-Wash Buffer (20mM HEPES, 150mM NaCl, 0.5mM Spermidine, 1x PIC, 0.05% Digitonin). 1200ng/mL of pA-MNase fusion protein (gift from Dr. Henikoff and Dr. K. Sarma) was added and beads were rotated at 4°C for 1 hr. Beads were washed twice then resuspended in Dig-Wash Buffer. Targeted digestion was performed at 0°C for 30 min by adding CaCl2 to 2mM. The digestion was stopped by adding one volume of 2X STOP Buffer (340mM NaCl, 20mM EDTA, 4mM EGTA, 0.02% Digitonin, 50ug/mL RNase A (Cat #3335399001, Roche), 50ug/mL Linear Acrylamide (Cat #K548, VWR), 2pg/mL Yeast Spike-In DNA). The target chromatin was released into the supernatant by shaking beads at 300rpm for 10 min at 37°C. The supernatant was then transferred to fresh tubes and incubated at 70°C for 10 min with SDS to 0.1% and 50ug of Proteinase K (Cat # EO0491, Thermo Scientific). DNA was then isolated with phenol:chloroform:isoamyl alcohol (Cat # P3803, Sigma-Aldrich) followed by precipitation with ammonium acetate. Libraries were prepared with 1.5ng or 5ng of CUT&RUN DNA using NEBNext Ultra II DNA Library Prep Kit for Illumina (Cat # E7645S, NEB) following the manufacturers protocol. CUT&RUN libraries were paired-end sequenced on a 300 cycle flow cell using either a NextSeq2000 or NovaSeq6000 system.

### Data Processing for CUT&RUN

Adapters were removed using Trimmomatic (V0.32, https://github.com/usadellab/Trimmomatic). For allele-specific analyses, SNPsplit (0.5.0)^53^ was used to generate an “N-masked” version of the mouse reference genome (mm10), wherein all SNPs between the *Mus musculus* C57BL/6 and *Mus musculus* CAST/EiJ mouse strains are masked by an “N” nucleotide. Reads were mapped to the “N-masked” genome using Bowtie2 (2.3.4.1) with the parameters [-q --local --very-sensitive-local --soft-clipped-unmapped-tlen --dovetail --no-mixed --no-discordant -- phred33 -I 10 -X 1000]^54^. For non-allele-specific analyses, reads were mapped to mm10 using Bowtie2 (2.3.4.1) with the same parameters as above. Duplicated reads were removed using Picard MarkDuplicates (1.141, https://broadinstitute.github.io/picard/) with option [REMOVE-DUPLICATES=true]. Reads mapped to regions from the ENCODE blacklist were removed. Allele-specific BAM files were then generated using SNPsplit (0.5.0) with option [--paired], wherein reads overlapping SNPs were tagged and sorted into Bl6 and Cast files. CPM normalized bigwig files were generated from allele-specific and non-allele-specific BAM files using deepTools bamCoverage (3.5.1) with options [--effectiveGenomeSize 2652783500 --normalizeUsing CPM -- ignoreForNormalization chrX]^55^. CPM normalized biological replicates were combined with WiggleTools mean (1.0)^56^ and visualized with Integrative Genomics Viewer (IGV, v2.14.0)^57^.

### Computational Analyses for CUT&RUN

#### d-score analysis

Promoter regions were defined as ±1kb from the TSS for all X-linked genes. Gene bodies were defined as 1kb downstream of the TSS to the TES. Gene bodies of X-linked genes shorter than 1kb were excluded from the analysis. Intergenic regions were defined as 10kb windows (generated using GenomicRanges (1.44.0)^58^) that did not overlap gene bodies or their promoter regions (1kb upstream of TSS to TES). Reads were assigned to defined windows using featureCounts from the Rsubread package (2.6.4)^59^ with arguments [useMetaFeatures = F, isPairedEnd = T]. For each biological replicate, d-scores [(reads^Bl6^/(reads^Cast^+ reads^Bl6^))-0.5] were calculated for promoters, gene bodies, and intergenic regions. Biological replicates were then averaged for each timepoint. P-values were calculated using a Wilcoxon signed-rank test with Benjamini Hochberg correction.

#### Heterochromatic Histone Mark Enrichment on the Xi

Reads from the *cast* files (allele with the Xi) were counted in 10kb windows across the genome (generated with GenomicRanges (1.44.0)^58^) using featureCounts from the Rsubread package (2.6.4)^59^ with arguments [useMetaFeatures = F, isPairedEnd = T]. Consensus autosomal peaks were then use to calculate a normalization factor using the trimmed mean of M-values (TMM) method from the edgeR package (3.34.1)^60^. To determine consensus autosomal peaks, peaks were first called on Cast files using Sparse Enrichment Analysis for CUT&RUN (SEACR)^61^ with options [norm relaxed] and then merged across all timepoints and biological replicates for each mark using bedtools intersect (2.29.1)^62^ with parameters [-nonamecheck -a]. To correct for chromatin accessibility or mappability bias, 10kb windows with a large variability in the IgG samples (read counts exceeded ±2 standard deviations from the mean) were discarded from the analysis. Read counts for each 10kb window were then averaged between biological replicates. Locally estimated scatterplot smoothing (LOESS) regression across all 10kb bins and Pearson correlation between each mark was performed.

#### Heterochromatic Histone Mark Accumulation Analyses

Visualization of heterochromatic mark accumulation across the Xi as compared to time 0 was performed as previously described^4^. Reads in 10kb bins on the *cast* allele were quantified and normalized using consensus autosomal peaks as described above. Read counts for each window were then averaged between biological replicates. Finally, to assess level of accumulation, normalized read counts from time 0h were subtracted from read counts at time 12h and 24h. Locally estimated scatterplot smoothing (LOESS) regression was performed on delta accumulation values across all 10kb bins.

#### Histone mark density across gene bodies

EnrichedHeatmap (1.22.0)^63^ was used to visualized hsitone mark density ±5kb of X-linked genes using CPM normalized bedgraphs and parameters [mean_mode = “w0”, w = 50, background = 0, smooth = T]. X-linked genes expressed in B cells were defined as those with an RPKM >1 in ≥2 biological replicates in previously published allele-specific RNA-seq data^26^. B cell XCI escape genes were previously reported in Sierra, I. *et. al.*^26^.

#### Temporal Analysis of Heterochromatic Histone Mark Accumulation

d-scores were calculated as described previously for gene bodies (TSS to TES) of all X-linked genes. Genes that accumulate marks with stimulation were determined by performing unsupervised k-means clustering on all X-linked genes and selecting clusters in which mark accumulated between time 0h and 24h. Early, Steady, and Late Accumulation groups were then assigned by performing k-means clustering for 3 temporal clusters on genes that accumulate mark. LINE, SINE, and CpG Island coordinates were downloaded from UCSC Table Browser.

#### Peak Differential Binding Analysis

Genome-wide differentially bound peaks on the Cast allele between 0h and 24h were determined for Ring1b and Suz12. Peaks were called on Cast files using Sparse Enrichment Analysis for CUT&RUN (SEACR)^61^ with options [0.01 non stringent] and analyzed for differential binding with DiffBind (3.2.7) in R.

#### Xist Deletion Analyses at Promoters, Gene Bodies, and Intergenic Regions

Promoters, gene bodies, and intergenic regions are defined as stated previously. Reads in these regions were quantified using featureCounts from the Rsubread package (2.6.4)^59^ with arguments [useMetaFeatures = F, isPairedEnd = T] and normalized using consensus autosomal peaks as described previously. For naïve samples Xist cKO samples (Figure 5), biological replicates were averaged and log2(Xist cKO/WT) was calculated. To determine defects in mark accumulation post stimulation (Figures 6 and **7**), log2 fold change between 0h and 24h was calculated for paired samples (in which 0h and 24h originated from the same mouse) then log2 fold change values were averaged within the WT and Xist cKO/TAT KO groups.

### Whole Genome Bisulfite Sequencing (WGBS)

Genomic DNA was extracted from B cells as follows: 2 million cells were resuspended in digestion buffer (100mM NaCl, 10mM Tris-HCl pH 8.0, 25mM EDTA pH 8.0, 0.5% SDS, 0.4mg/mL Proteinase K (Cat # EO0491, Thermo Scientific)) and incubated at 56°C for 2 hrs. DNA was extracted with an equal volume of phenol:chloroform:isoamyl alcohol (Cat # P3803, Sigma-Aldrich) followed by precipitation with ammonium acetate. Bisulfite libraries were prepared using the NEBNext Ultra II DNA Library Prep Kit for Illumina (Cat # E7645S, NEB) as follows: 1.25 ug of gDNA was spiked with 0.1% Unmethylated Lambda DNA (D152A, Promega) and fragmented with a Covaris S220 using parameters (PIP: 175, DF: 10%, C/B: 200, Temp: 7°C, Time: 180 seconds). DNA was then end repaired and ligated to the NEBNext Methylated Adaptor for Illumina (Cat # E7535, NEB). DNA was bisulfite converted using the EZ DNA Methylation-Gold Kit (Cat # D5005, Zymo Research) following manufacturers protocol. Bisulfite converted DNA was then amplified for 8 cycles using Q5U Hot Start High-Fidelity DNA Polymerase (Cat # M0515S, NEB). Libraries were paired-end sequenced on a P3 300 flow cell using the NextSeq2000 system.

### Data Processing for Allele-Specific WGBS

Adapters were removed using Trimmomatic (V0.32, https://github.com/usadellab/Trimmomatic). SNPsplit (0.5.0)^53^ was used to generate an “N-masked” version of the mouse reference genome (mm10) as detailed above. Reads were mapped to the “N-masked” genome using Bismark (0.22.3) with the parameters [-- un --nucleotide_coverage --ambiguous]^64^. Non-bisulfite converted reads were removed using Bismark filter_non_conversion (0.23.1) with parameters [-p --minimum_count 5 -- percentage_cutoff 50] and duplicates were removed using Bismark deduplicate_bismark (0.23.1). Allele-specific BAM files were then generated using SNPsplit (0.5.0) with options [--paired --bisulfite]. Methylation calls were extracted using Bismark bismark_methylation_extractor (0.23.1) with parameters [--ignore 4 --ignore_r2 10 -p --bedGraph --gzip]. Biological replicates for each timepoint were combined using methylKit unite (1.18.0)^65^.

### Computational Analysis for Allele-Specific WGBS

Promoter regions were defined as ±1kb from the TSS for all X-linked genes. CpG islands coordinates for mm10 were downloaded from UCSC Table Browser. Percent DNA methylation at promoter regions and CpG islands was quantified using methylKit regionCounts (1.18.0) and methylKit percMethylation (1.18.0). To plot gene body DNA methylation ±5kb of X-linked genes, percent DNA methylation in 500bp windows across the genome was first quantified using methylKit tileMethylCounts (1.18.0) with arguments [win.size = 500, step.size = 500, cov.bases = 0] and methylKit percMethylation (1.18.0). This was then visualized using EnrichedHeatmap (1.22.0)^63^ with parameters [extend = 5000, mean_mode = “absolute”, w = 50, background =NA].

### RNA Isolation and qPCR

Total RNA was extracted using TRIzol Reagent (Cat # 15596018, Thermo Fisher Scientific). 500ng of RNA was converted to cDNA using qScript cDNA SuperMix (Cat # 95048-025, QuantaBio). To quantify Xist RNA levels, PerfeCTa SYBR GreenSupermix, Low ROX (Cat # 95056-500, QuantaBio) and a QuantStudio 6 FlexSystem (Applied Biosystems) were used.

### Western Blot

Cells were harvested and washed twice with ice cold phosphate buffered saline (PBS; Cat # P5368, Sigma-Aldrich). Cells were resuspended in Triton Extraction Buffer (TEB; PBS containing 0.5% Triton X 100 (v/v), 2 mM phenylmethylsulfonyl fluoride (PMSF), 0.02% (w/v) NaN3) and lysed on ice for 10 minutes. Tubes were centrifuged at 650 x g for 10 min at 4°C to spin down the nuclei. Nuclei were washed with TEB, pelleted at 650 x g for 10 min at 4°C, then resuspended in 0.2 N HCl at a density of 4x107 nuclei per ml. Histones were acid extracted overnight at 4°C. Samples were centrifuged at 650 x g for 10 min at 4°C to pellet debris, supernatant was transferred to a clean tube, and 1/10 the volume of 2M NaOH was added. Protein concentration was determined using the Pierce BCA Protein Assay Kit (Cat # 23225, Thermo Scientific). 3.25ug of protein was resolved on an SDS-PAGE gel (Cat # NP0335, Invitrogen), transferred to a nitrocellulose membrane (Cat # 88018, Thermo Scientific), and Ponceau stained for 10 minutes at room temperature (Cat # A40000279, Thermo Scientific). Following imaging, membranes were washed, blocked for 1 hour at room temperature, then incubated with the H3K27me3 primary antibody at 4°C overnight (1:1000; Cat #39055, Active Motif). Membranes were washed, incubated for 1 hour at room temperature with HRP-conjugated secondary antibody (Cat # ab6721, Abcam), and visualized using Pierce ECL Western Substrate (Cat # 32106, Thermo Scientific) and a ChemiDoc Imaging system (BioRad).

## REFERENCES

1 Lyon, M. F. Gene Action in the X-chromosome of the Mouse (Mus musculus L.). Nature 190, 372–373 (1961). 10.1038/190372a0

2 Brown, C. J. et al. The human XIST gene: Analysis of a 17 kb inactive X-specific RNA that contains conserved repeats and is highly localized within the nucleus. Cell 71, 527–542 (1992). 10.1016/0092-8674(92)90520-M

3 Brockdorff, N. et al. The product of the mouse Xist gene is a 15 kb inactive X-specific transcript containing no conserved ORF and located in the nucleus. Cell 71, 515–526 (1992). 10.1016/0092-8674(92)90519-i

4 Żylicz, J. J., et al. The Implication of Early Chromatin Changes in X Chromosome Inactivation. Cell 176, 182–197.e123 (2019). 10.1016/j.cell.2018.11.041

5 McHugh, C. A. et al. The Xist lncRNA interacts directly with SHARP to silence transcription through HDAC3. Nature 521, 232–236 (2015). 10.1038/nature14443

6 Silva, J. et al. Establishment of histone h3 methylation on the inactive X chromosome requires transient recruitment of Eed-Enx1 polycomb group complexes. Dev Cell 4, 481–495 (2003). 10.1016/s1534-5807(03)00068-6

7 Plath, K. et al. Role of histone H3 lysine 27 methylation in X inactivation. Science 300, 131–135 (2003). 10.1126/science.1084274

8 Engreitz, J. M. et al. The Xist lncRNA exploits three-dimensional genome architecture to spread across the X chromosome. Science 341, 1–9 (2013). 10.1126/science.1237973

9 de Napoles, M. et al. Polycomb group proteins Ring1A/B link ubiquitylation of histone H2A to heritable gene silencing and X inactivation. Dev Cell 7, 663–676 (2004). 10.1016/j.devcel.2004.10.005

10 Keniry, A. et al. Setdb1-mediated H3K9 methylation is enriched on the inactive X and plays a role in its epigenetic silencing. Epigenetics and Chromatin 9, 1–20 (2016). 10.1186/s13072-016-0064-6

11 Ichihara, S., Nagao, K., Sakaguchi, T., Obuse, C. & Sado, T. SmcHD1 underlies the formation of H3K9me3 blocks on the inactive X chromosome in mice. *Development (Cambridge*, England) 149, 1–30 (2022). 10.1242/dev.200864

12 Norris, D. P., Brockdorff, N. & Rastan, S. Methylation status of CpG-rich islands on active and inactive mouse X chromosomes. Mammalian Genome 1, 78–83 (1991). 10.1007/BF02443782

13 Lock, L. F., Takagi, N. & Martin, G. R. Methylation of the Hprt gene on the inactive X occurs after chromosome inactivation. Cell 48, 39–46 (1987). 10.1016/0092-8674(87)90353-9

14 Gendrel, A.-V. et al. Smchd1-Dependent and -Independent Pathways Determine Developmental Dynamics of CpG Island Methylation on the Inactive X Chromosome. Developmental Cell 23, 265–279 (2012). 10.1016/j.devcel.2012.06.011

15 Simon, M. D. et al. High-resolution Xist binding maps reveal two-step spreading during X-chromosome inactivation. Nature 504, 465–469 (2013). 10.1038/nature12719

16 Colognori, D., Sunwoo, H., Kriz, A. J., Wang, C. Y. & Lee, J. T. Xist Deletional Analysis Reveals an Interdependency between Xist RNA and Polycomb Complexes for Spreading along the Inactive X. Molecular Cell 74, 101–117.e110 (2019). 10.1016/j.molcel.2019.01.015

17 Almeida, M. et al. PCGF3/5-PRC1 initiates Polycomb recruitment in X chromosome inactivation. Science 356, 1081–1084 (2017). 10.1126/science.aal2512

18 da Rocha, S. T. et al. Jarid2 Is Implicated in the Initial Xist-Induced Targeting of PRC2 to the Inactive X Chromosome. Mol Cell 53, 301–316 (2014). 10.1016/j.molcel.2014.01.002

19 Cooper, S. et al. Jarid2 binds mono-ubiquitylated H2A lysine 119 to mediate crosstalk between Polycomb complexes PRC1 and PRC2. Nat Commun 7, 13661 (2016). 10.1038/ncomms13661

20 Berletch, J. B. et al. Escape from X Inactivation Varies in Mouse Tissues. PLoS Genetics 11, 1–26 (2015). 10.1371/journal.pgen.1005079

21 Tukiainen, T. et al. Landscape of X chromosome inactivation across human tissues. Nature 550, 244–248 (2017). 10.1038/nature24265

22 Cotton, A. M., Price, E. M., Jones, M. J., Balaton, B. P., Kobor, M. S. & Brown, C. J. Landscape of DNA methylation on the X chromosome reflects CpG density, functional chromatin state and X-chromosome inactivation. Human Molecular Genetics 24, 1528–1539 (2015). 10.1093/hmg/ddu564

23 Wang, J., Syrett, C. M., Kramer, M. C., Basu, A., Atchison, M. L. & Anguera, M. C. Unusual maintenance of X chromosome inactivation predisposes female lymphocytes for increased expression from the inactive X. Proceedings of the National Academy of Sciences 113, E2029–E2038 (2016). 10.1073/pnas.1520113113

24 Syrett, C. M. et al. Loss of Xist RNA from the inactive X during B cell development is restored in a dynamic YY1-dependent two-step process in activated B cells. PLOS Genetics 13, e1007050–e1007050 (2017). 10.1371/journal.pgen.1007050

25 Pyfrom, S. et al. The dynamic epigenetic regulation of the inactive X chromosome in healthy human B cells is dysregulated in lupus patients. Proceedings of the National Academy of Sciences of the United States of America 118 (2021). 10.1073/pnas.2024624118

26 Sierra, I. et al. Remodeling and compaction of the inactive X is regulated by Xist during female B cell activation. (2022). 10.1101/2022.10.19.512821

27 Kong, W. et al. Increased expression of Bruton’s tyrosine kinase in peripheral blood is associated with lupus nephritis. Clin Rheumatol 37, 43–49 (2018). 10.1007/s10067-017-3717-3

28 Forsyth, K. S., Jiwrajka, N., Lovell, C. D., Toothacre, N. E. & Anguera, M. C. The conneXion between sex and immune responses. Nature Reviews Immunology 24, 487–502 (2024). 10.1038/s41577-024-00996-9

29 Hellman, A. & Chess, A. Gene body-specific methylation on the active X chromosome. Science 315, 1141–1143 (2007). 10.1126/science.1136352

30 Baubec, T. et al. Genomic profiling of DNA methyltransferases reveals a role for DNMT3B in genic methylation. Nature 520, 243–U278 (2015). 10.1038/nature14176

31 Balaton, B. P. & Brown, C. J. Contribution of genetic and epigenetic changes to escape from X-chromosome inactivation. Epigenet Chromatin 14 (2021). ARTN 30 10.1186/s13072-021-00404-9

32 Ku, M. et al. Genomewide analysis of PRC1 and PRC2 occupancy identifies two classes of bivalent domains. PLoS Genet 4, e1000242 (2008). 10.1371/journal.pgen.1000242

33 Li, H. et al. Polycomb-like proteins link the PRC2 complex to CpG islands. Nature 549, 287–291 (2017). 10.1038/nature23881

34 Knutson, S. K. et al. Durable tumor regression in genetically altered malignant rhabdoid tumors by inhibition of methyltransferase EZH2. Proceedings of the National Academy of Sciences of the United States of America 110, 7922–7927 (2013). 10.1073/pnas.1303800110

35 Sasaki, T., Katagi, H., Goldman, S., Becher, O. J. & Hashizume, R. Convection-Enhanced Delivery of Enhancer of Zeste Homolog-2 (EZH2) Inhibitor for the Treatment of Diffuse Intrinsic Pontine Glioma. Neurosurgery 87, E680–E688 (2020). 10.1093/neuros/nyaa301

36 Guo, M. et al. EZH2 Represses the B Cell Transcriptional Program and Regulates Antibody-Secreting Cell Metabolism and Antibody Production. Journal of Immunology 200, 1039–1052 (2018). 10.4049/jimmunol.1701470

37 Hobeika, E. et al. Testing gene function early in the B cell lineage in mb1-cre mice. Proceedings of the National Academy of Sciences of the United States of America 103, 13789–13794 (2006). 10.1073/pnas.0605944103

38 Lovell, C. D., Jiwrajka, N., Amerman, H. K., Cancro, M. P. & Anguera, M. C. Xist Deletion in B Cells Results in Systemic Lupus Erythematosus Phenotypes. bioRxiv (2024). 10.1101/2024.05.15.594175

39 Joshi, S. K., Hashimoto, K. & Koni, P. A. Induced DNA recombination by Cre recombinase protein transduction. Genesis 33, 48–54 (2002). 10.1002/gene.10089

40 Zaprazna, K. & Atchison, M. L. YY1 Controls Immunoglobulin Class Switch Recombination and Nuclear Activation-Induced Deaminase Levels. Molecular and Cellular Biology 32, 1542–1554 (2012). 10.1128/Mcb.05989-11

41 Pintacuda, G. et al. hnRNPK Recruits PCGF3/5-PRC1 to the Xist RNA B-Repeat to Establish Polycomb-Mediated Chromosomal Silencing. Mol Cell 68, 955–969 e910 (2017). 10.1016/j.molcel.2017.11.013

42 Lee, Y. W., Weissbein, U., Blum, R. & Lee, J. T. G-quadruplex folding in Xist RNA antagonizes PRC2 activity for stepwise regulation of X chromosome inactivation. Molecular Cell 84 (2024). 10.1016/j.molcel.2024.04.015

43 Forsyth, K. S. et al. Maintenance of X chromosome inactivation after T cell activation requires NF-kappaB signaling. Sci Immunol 9, eado0398 (2024). 10.1126/sciimmunol.ado0398

44 Chadwick, B. P. & Willard, H. F. Multiple spatially distinct types of facultative heterochromatin on the human inactive X chromosome. Proceedings of the National Academy of Sciences of the United States of America 101, 17450–17455 (2004). 10.1073/pnas.0408021101

45 Rens, W., Wallduck, M. S., Lovell, F. L., Ferguson-Smith, M. A. & Ferguson-Smith, A. C. Epigenetic modifications on X chromosomes in marsupial and monotreme mammals and implications for evolution of dosage compensation. Proceedings of the National Academy of Sciences of the United States of America 107, 17657–17662 (2010). 10.1073/pnas.0910322107

46 Suzuki, M. M. & Bird, A. DNA methylation landscapes: provocative insights from epigenomics. Nature Reviews Genetics 9, 465–476 (2008). 10.1038/nrg2341

47 Shen, X. H. et al. EZH1 Mediates Methylation on Histone H3 Lysine 27 and Complements EZH2 in Maintaining Stem Cell Identity and Executing Pluripotency. Molecular Cell 32, 491–502 (2008). 10.1016/j.molcel.2008.10.016

48 Li, W. et al. Targeting EZH1/2 induces cell cycle arrest and inhibits cell proliferation through reactivation of p57 and TP53INP1 in mantle cell lymphoma. Cancer Biol Med 16, 530–541 (2019). 10.20892/j.issn.2095-3941.2018.0380

49 Xu, C. et al. Binding of different histone marks differentially regulates the activity and specificity of polycomb repressive complex 2 (PRC2). Proceedings of the National Academy of Sciences of the United States of America 107, 19266–19271 (2010). 10.1073/pnas.1008937107

50 Cao, R. et al. Role of histone H3 lysine 27 methylation in polycomb-group silencing. Science 298, 1039–1043 (2002). 10.1126/science.1076997

51 Kuzmichev, A., Nishioka, K., Erdjument-Bromage, H., Tempst, P. & Reinberg, D. Histone methyltransferase activity associated with a human multiprotein complex containing the Enhancer of Zeste protein. Gene Dev 16, 2893–2905 (2002). 10.1101/gad.1035902

52 Skene, P. J. & Henikoff, S. An efficient targeted nuclease strategy for high-resolution mapping of DNA binding sites. Elife 6 (2017). 10.7554/eLife.21856

53 Krueger, F. & Andrews, S. R. SNPsplit: Allele-specific splitting of alignments between genomes with known SNP genotypes. F1000Research 5, 1479–1479 (2016). 10.12688/f1000research.9037.1

54 Langmead, B. & Salzberg, S. L. Fast gapped-read alignment with Bowtie 2. Nat Methods 9, 357–359 (2012). 10.1038/nmeth.1923

55 Ramirez, F. et al. deepTools2: a next generation web server for deep-sequencing data analysis. Nucleic Acids Res 44, W160–165 (2016). 10.1093/nar/gkw257

56 Zerbino, D. R., Johnson, N., Juettemann, T., Wilder, S. P. & Flicek, P. WiggleTools: parallel processing of large collections of genome-wide datasets for visualization and statistical analysis. Bioinformatics 30, 1008–1009 (2014). 10.1093/bioinformatics/btt737

57 Thorvaldsdottir, H., Robinson, J. T. & Mesirov, J. P. Integrative Genomics Viewer (IGV): high-performance genomics data visualization and exploration. Brief Bioinform 14, 178–192 (2013). 10.1093/bib/bbs017

58 Lawrence, M. et al. Software for computing and annotating genomic ranges. PLoS Comput Biol 9, e1003118 (2013). 10.1371/journal.pcbi.1003118

59 Liao, Y., Smyth, G. K. & Shi, W. The R package Rsubread is easier, faster, cheaper and better for alignment and quantification of RNA sequencing reads. Nucleic Acids Res 47, e47 (2019). 10.1093/nar/gkz114

60 Robinson, M. D., McCarthy, D. J. & Smyth, G. K. edgeR: a Bioconductor package for differential expression analysis of digital gene expression data. Bioinformatics 26, 139–140 (2010). 10.1093/bioinformatics/btp616

61 Meers, M. P., Tenenbaum, D. & Henikoff, S. Peak calling by Sparse Enrichment Analysis for CUT&RUN chromatin profiling. Epigenetics Chromatin 12, 42 (2019). 10.1186/s13072-019-0287-4

62 Quinlan, A. R. & Hall, I. M. BEDTools: a flexible suite of utilities for comparing genomic features. Bioinformatics 26, 841–842 (2010). 10.1093/bioinformatics/btq033

63 Gu, Z., Eils, R., Schlesner, M. & Ishaque, N. EnrichedHeatmap: an R/Bioconductor package for comprehensive visualization of genomic signal associations. BMC Genomics 19, 234 (2018). 10.1186/s12864-018-4625-x

64 Krueger, F. & Andrews, S. R. Bismark: a flexible aligner and methylation caller for Bisulfite-Seq applications. Bioinformatics 27, 1571–1572 (2011). 10.1093/bioinformatics/btr167

65 Akalin, A. et al. methylKit: a comprehensive R package for the analysis of genome-wide DNA methylation profiles. Genome Biology 13 (2012). 10.1186/gb-2012-13-10-r87

