## Supplementary figures and images for "Xist RNA Dependent and Independent Mechanisms Regulate Dynamic X Chromosome Inactivation in B Lymphocytes"

### Supplemental Figures

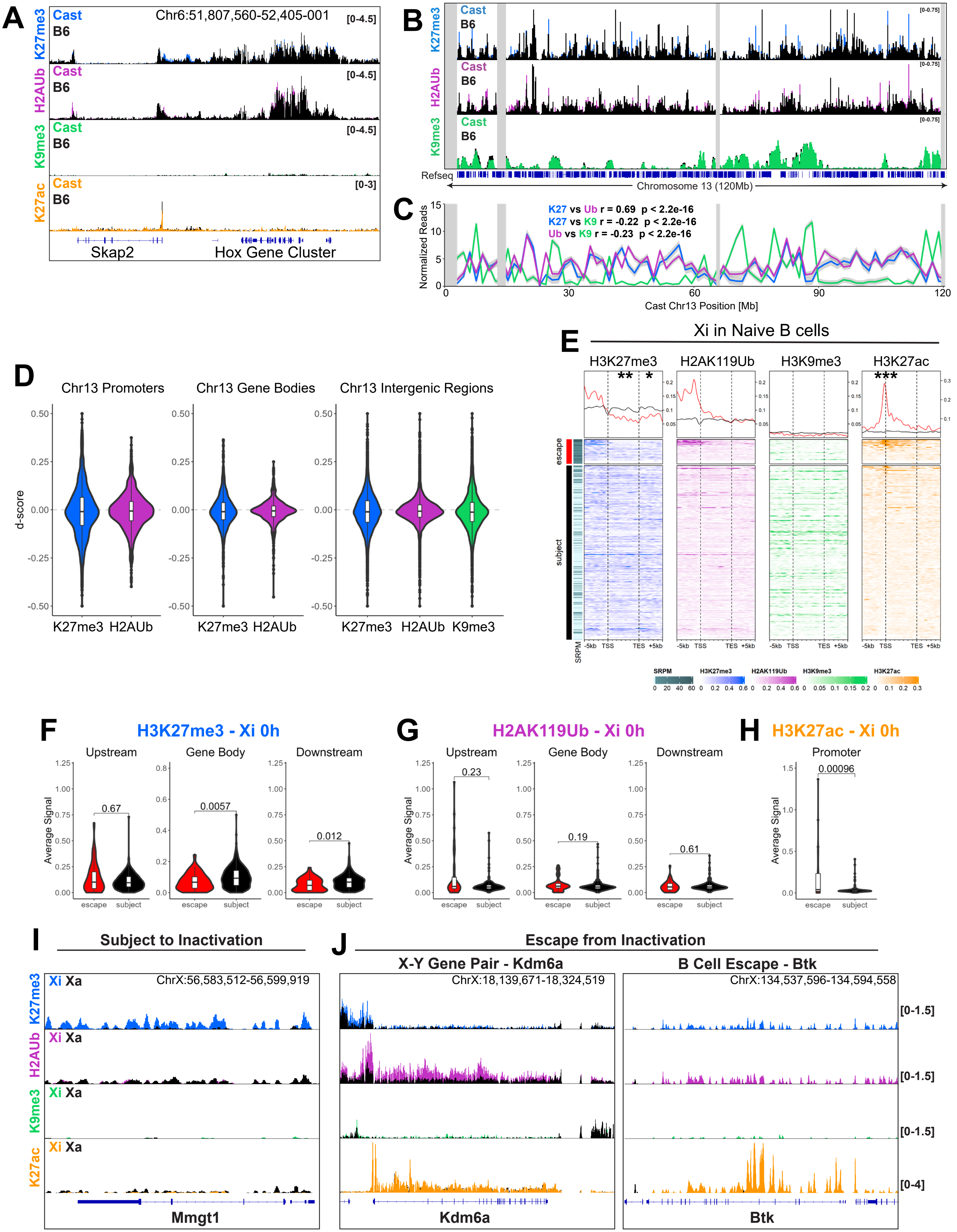

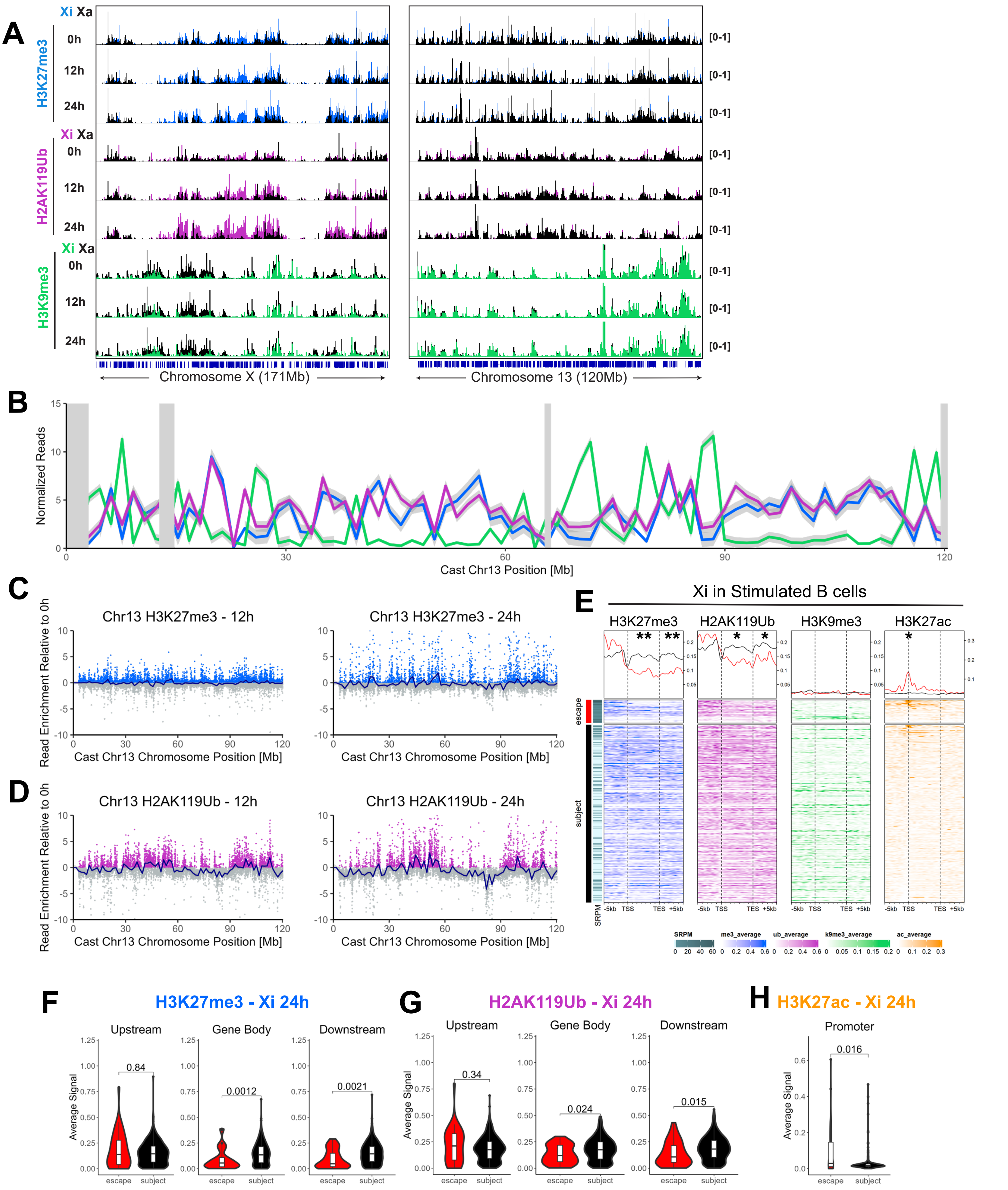

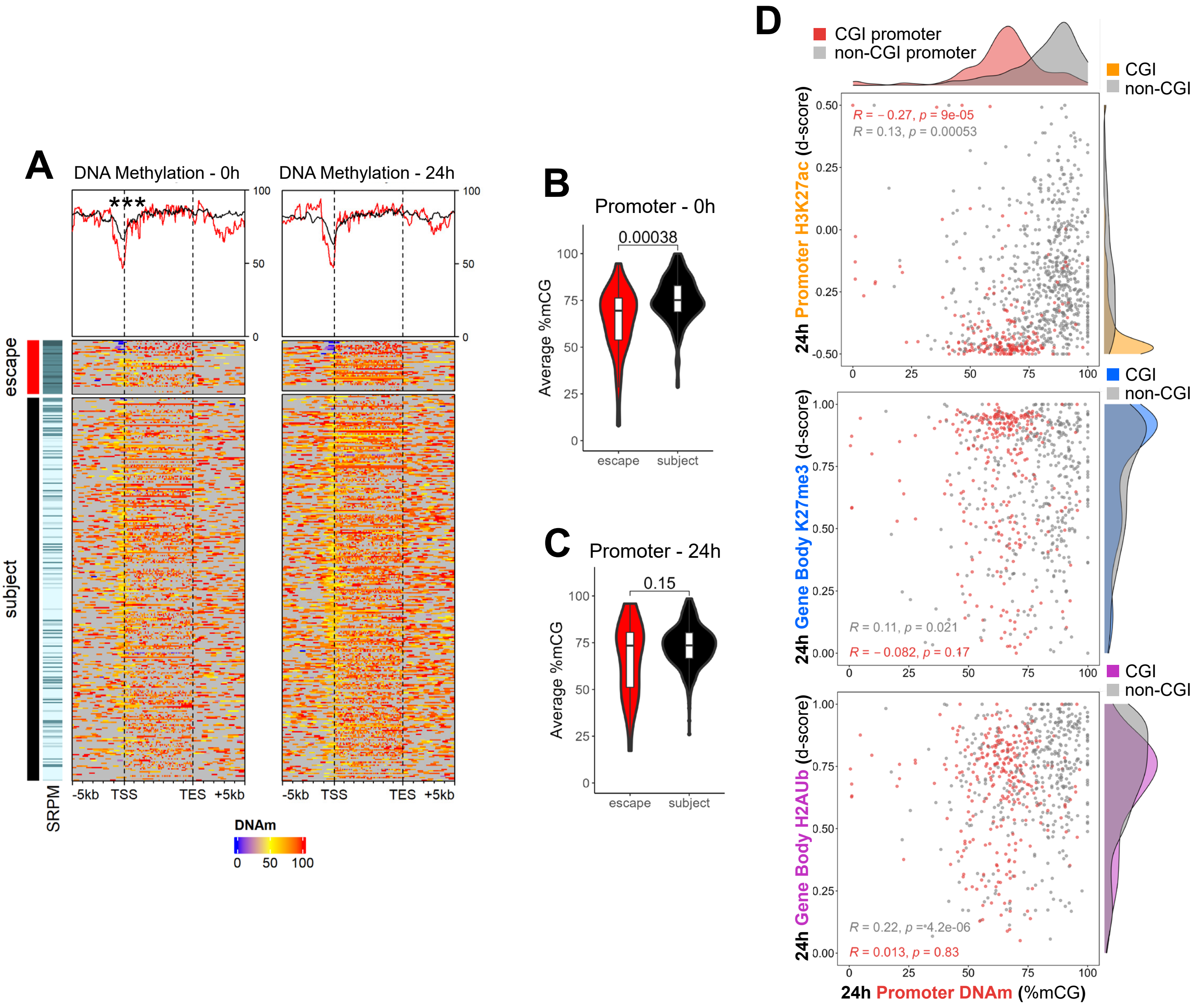

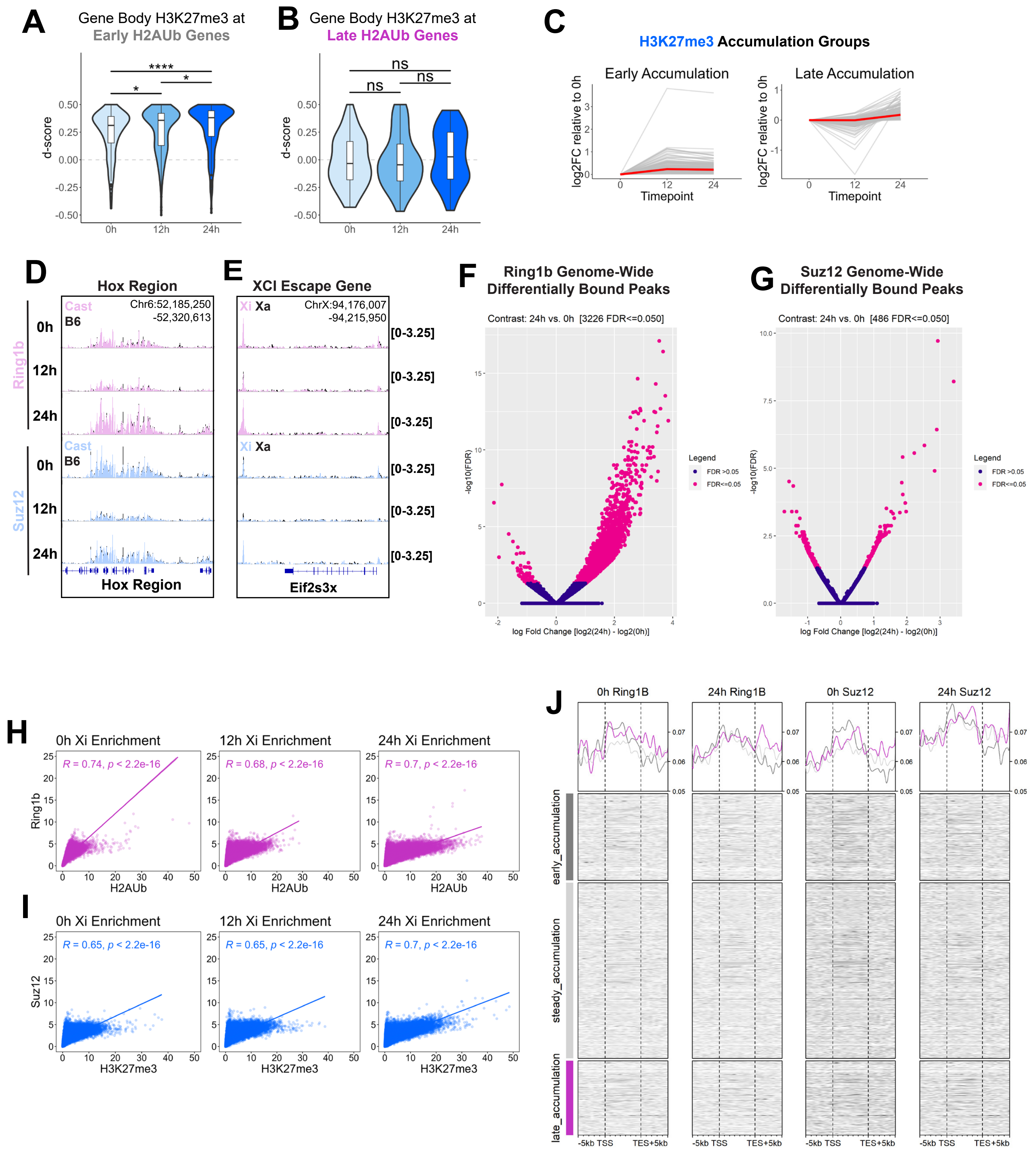

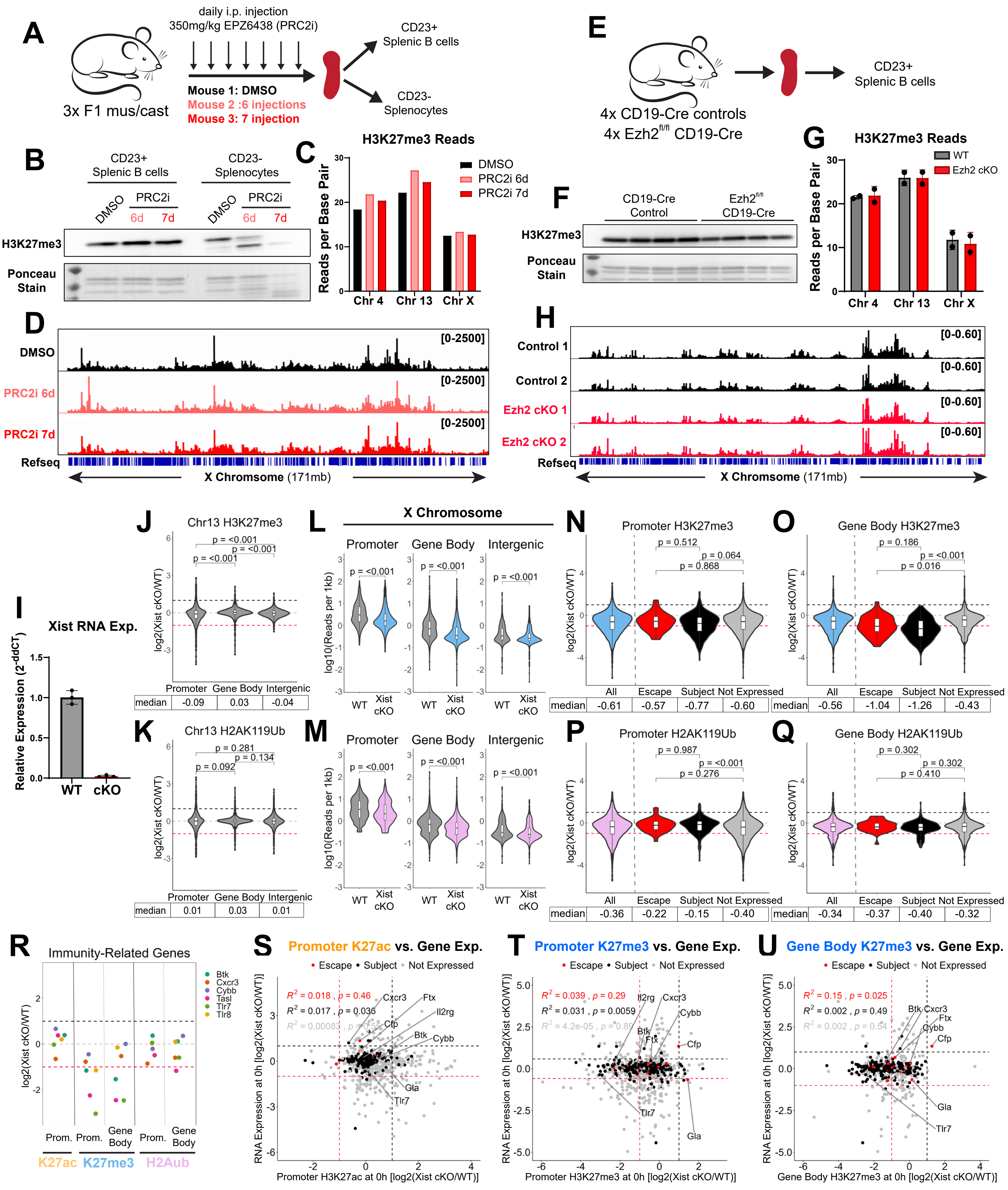

#### Figure S6

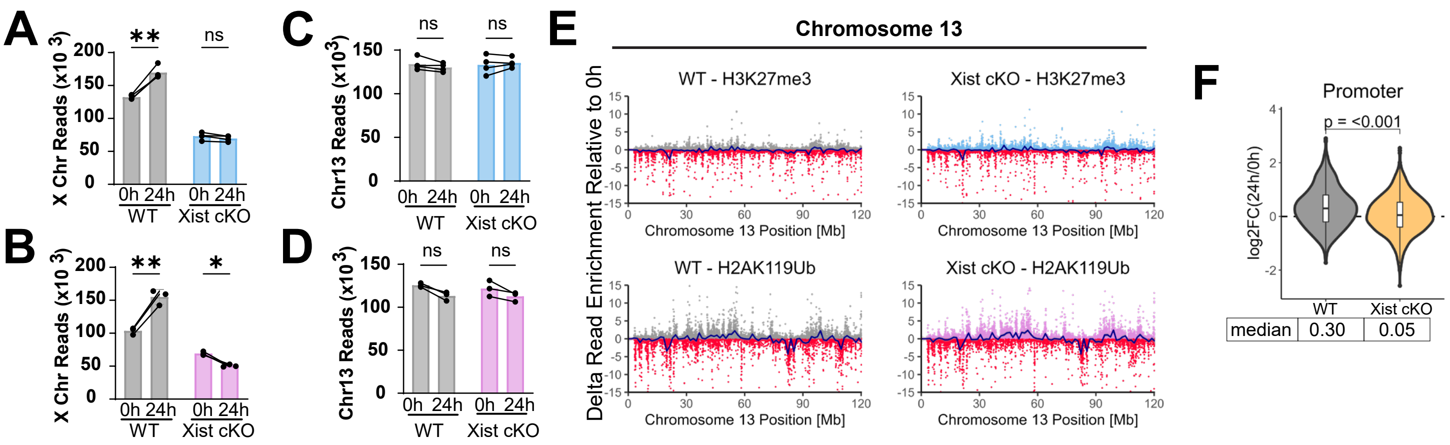


| median | 0.30 | 0.05 |
|---|---|---|

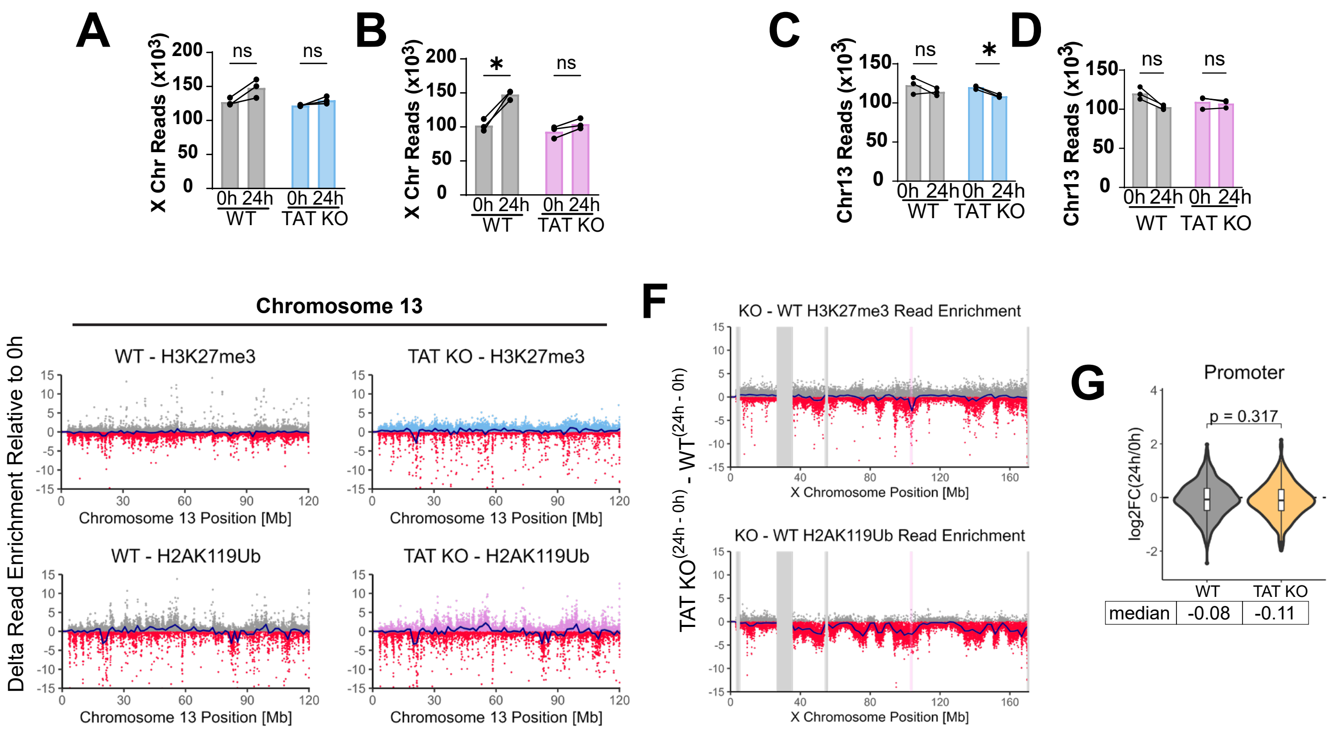
